# Long Wake/Short Sleep Bouts and Hyperactivity with Advanced Age in a Mouse Model of Early Onset Alzheimer’s Disease

**DOI:** 10.64898/2025.12.29.696818

**Authors:** Ryan K. Tisdale, Yu Sun, Stephanie R. Miller, Stephanie M. Lee, Sunmee Park, Jia Shin, Giancarlo Allocca, Jorge J. Palop, Thomas S. Kilduff

## Abstract

Poor sleep quality and reduced sleep duration are associated with Alzheimer’s disease (AD)-related β-amyloid (Aβ) pathologies. We conducted two studies of sleep/wake, activity and body temperature in *App*^NL-G-F^ mice, a strain that exhibits three mutations in the human *App* gene associated with elevated risk for early onset AD. First, *App*^NL-G-F^ mice were compared to wildtype (WT) littermates at 14-18 and 18-22 months of age and, at both ages, were found to exhibit more Wake and less NREM and REM sleep than WT littermates. This long wake/short sleep phenotype was evident during the dark phase at 14-18 months but occurred in both the light and dark phases at 18-22 months. *App*^NL-G-F^ mice had fewer short (<60 sec) and more long (>260 sec) Wake bouts and were hyperactive at 18-22 months, which undoubtedly contributed to the increased Wake/reduced sleep. Despite this reduced sleep phenotype, *App*^NL-G-F^ mice were no sleepier than WT mice and the sleep homeostat was functional in both strains. In the second study, sex differences in these parameters were assessed at 18-24 months. Reduced sleep was evident in both sexes of *App*^NL-G-F^ mice but was clearly more evident in females. Wake and REM sleep bout durations were longer in both sexes of *App*^NL-G-F^ mice than in WT littermates. EEG spectral power during NREM sleep was reduced in female *App*^NL-G-F^ mice between 4.88-10.50 Hz compared to WT mice whereas, during REM sleep, both male and female *App*^NL-G-F^ mice exhibited reduced spectral power in the theta range. These results suggest that Aβ deposition may impair state transition mechanism(s) in *App*^NL-G-F^ mice and demonstrate that, as in human AD patients, the long wake/short sleep phenotype was more evident in female *App*^NL-G-F^ mice, thus supporting the use of this strain as a model to investigate interventions that mitigate AD burden during early disease stages.

## Introduction

Alzheimer’s disease (AD) is a progressive neurodegenerative disease with a preclinical/prodromal stage characterized by extracellular accumulation of amyloid-β (Aβ) into senile Aβ plaques and later by intracellular accumulation of hyperphosphorylated tau (hTau) protein into neurofibrillary tangles (NFTs), both of which contribute to neuronal and synaptic loss and clinical manifestations of cognitive dysfunction and memory impairment (Bateman et al., 2012; Jack et al., 2013; Vos et al., 2013). Aβ and hTau are byproducts of normal cellular processes and are cleaved from the *Amyloid Precursor Protein* (*App*) and the *Microtubule Associated Protein* (*MAPT*), respectively. While *MAPT* mutations are more common in Frontotemporal Dementia, several missense *App* mutations have been associated with elevated risk for familial or Early-Onset Alzheimer’s Disease (EOAD) (Carter et al., 1992; Chartier-Harlin et al., 1991; Clark et al., 1998; Goate et al., 1991; Hutton et al., 1998).

AD and sleep exhibit a bidirectional relationship whereby disordered sleep is both a symptom of AD pathologies and has a causative role in their development (Lucey et al., 2019; Musiek & Ju, 2022; Wang & Holtzman, 2020). Although disordered sleep occurs across all disease phases in AD, due to the causal role that disordered sleep may play in disease progression at the prodromal stage, considerable translational interest exists in understanding the mechanisms by which disordered sleep contributes to disease progression and the potential beneficial impacts of sleep-related interventions during the preclinical phase (Kang et al., 2009; Musiek & Ju, 2022; Roh et al., 2012). Sex differences have been documented in AD-related sleep dysfunction (C. E. Johnson et al., 2024) but the role that biological sex plays in determining AD risk is difficult to disentangle from impacts of sex on other biological processes and socioeconomic risk factors (Gong et al., 2023).

Sleep/wake has been characterized in several of the mouse models of AD that have been developed in the last two decades (Katsuki et al., 2022). Several amyloid and tau mutant mouse strains have been shown to exhibit sleep disruption and increased wakefulness consistent with human AD (Holth et al., 2017; Huitron-Resendiz et al., 2002; Jyoti et al., 2015; Kang et al., 2009; Platt et al., 2011; Roh et al., 2012; Schneider et al., 2014; Wisor et al., 2005; Bin Zhang et al., 2005; Zhu et al., 2018). Furthermore, chronic sleep restriction compounds pathology in amyloid and tau mutant mice compared to free-sleeping mutants (Kang et al., 2009; Zhu et al., 2018), whereas immunizing an AD model against Aβ appears to normalize disturbed sleep and daily Aβ cycling (Roh et al., 2012). Together, these studies support the concept of a bidirectional relationship between sleep/wake and the neuropathological components of AD. In the current study, we characterized sleep/wake states in aged *App*^NL-G-F^ mice, a murine model that exhibits pronounced amyloid pathology and which was engineered to exhibit three missense mutations in the *App* gene associated with an elevated risk for EOAD (Manabe & Saito, 2025; Saito et al., 2014). Although previous studies have characterized sleep/wake in 6-12 month old *App*^NL-G-F^ mice (Calafate et al., 2023; Maezono et al., 2020; Yao et al., 2023), we report results here from (1) a longitudinal comparison of *App*^NL-G-F^ and WT mice at 14-18 vs. 18-22 months of age, and (2) continuous 14-day EEG/EMG recordings of male vs. female *App*^NL-G-F^ and WT mice at 18-24 months of age when the pathology has become more severe. We find that *App*^NL-G-F^ mice exhibit an insomnia-like long wake/short sleep phenotype that is particularly pronounced in females and is suggestive of a defect in the mechanisms underlying transitions between arousal states.

## Results

### App^NL-G-F^ mice exhibit region-specific amyloid pathology and microglial activation

The design for the two experiments conducted in male and female *App*^NL-G-F^ mice is illustrated in **Figure 1** and described in detail in Materials and Methods. As described previously (Miller et al., 2024; Saito et al., 2014; Sasaguri et al., 2017), *App*^NL-G-F^ mice are characterized by progressive amyloidosis and gliosis and may thus model preclinical stages of AD. Amyloid deposition in representative 22 month old WT and *App*^NL-G-F^ mice is presented in **Figures 2A and 2A’**. Two-way ANOVA revealed a significant main effect of genotype on Aβ expression (*p* < 0.0005). **Figure 2A**” presents, for each of 5 brain areas, the proportion of each brain area in which Aβ expression was observed. *Post hoc* analysis indicated that Aβ expression was significantly higher in the retrospenial cortex (RSC) of *App*^NL-G-F^ mice compared to WT. Similarly, **Figure 2B and 2B’** compares Iba1 staining, an indicator of gliosis, between the two strains and **Figure 2B”** presents, for each of 5 brain areas, the proportion of that area in which Iba1 is expressed relative to WT mice. Two-way ANOVA also demonstrated a significant main effect of genotype for Iba1 expression (*p* < 0.0001); *post hoc* analysis revealed significantly higher Iba1 levels in *App*^NL-G-F^ mice relative to WT in all 5 brain regions examined.

**Figure 1.**
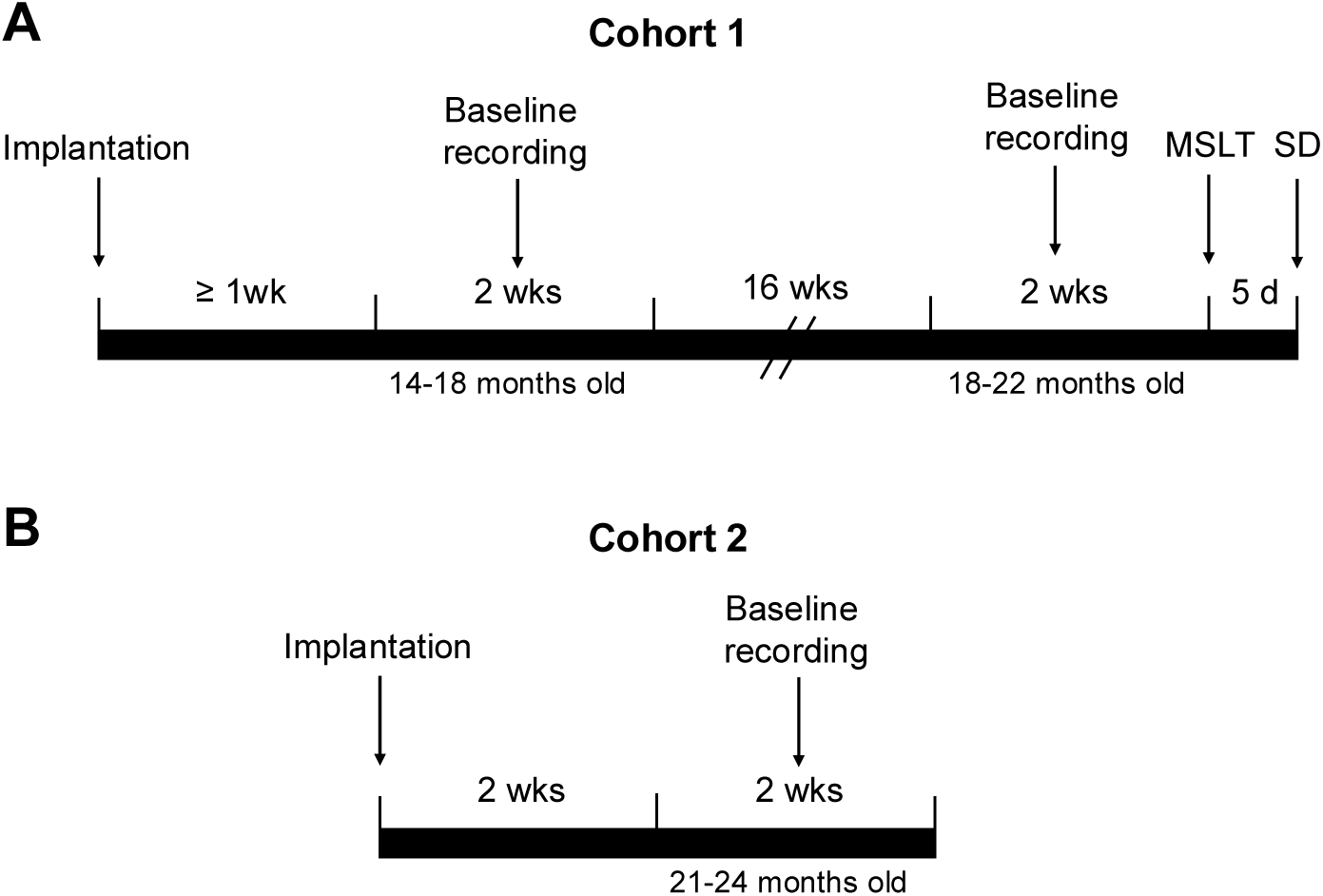
Timelines for the experimental procedures for (A) Cohort 1 and (B) Cohort 2.

**Figure 2.**
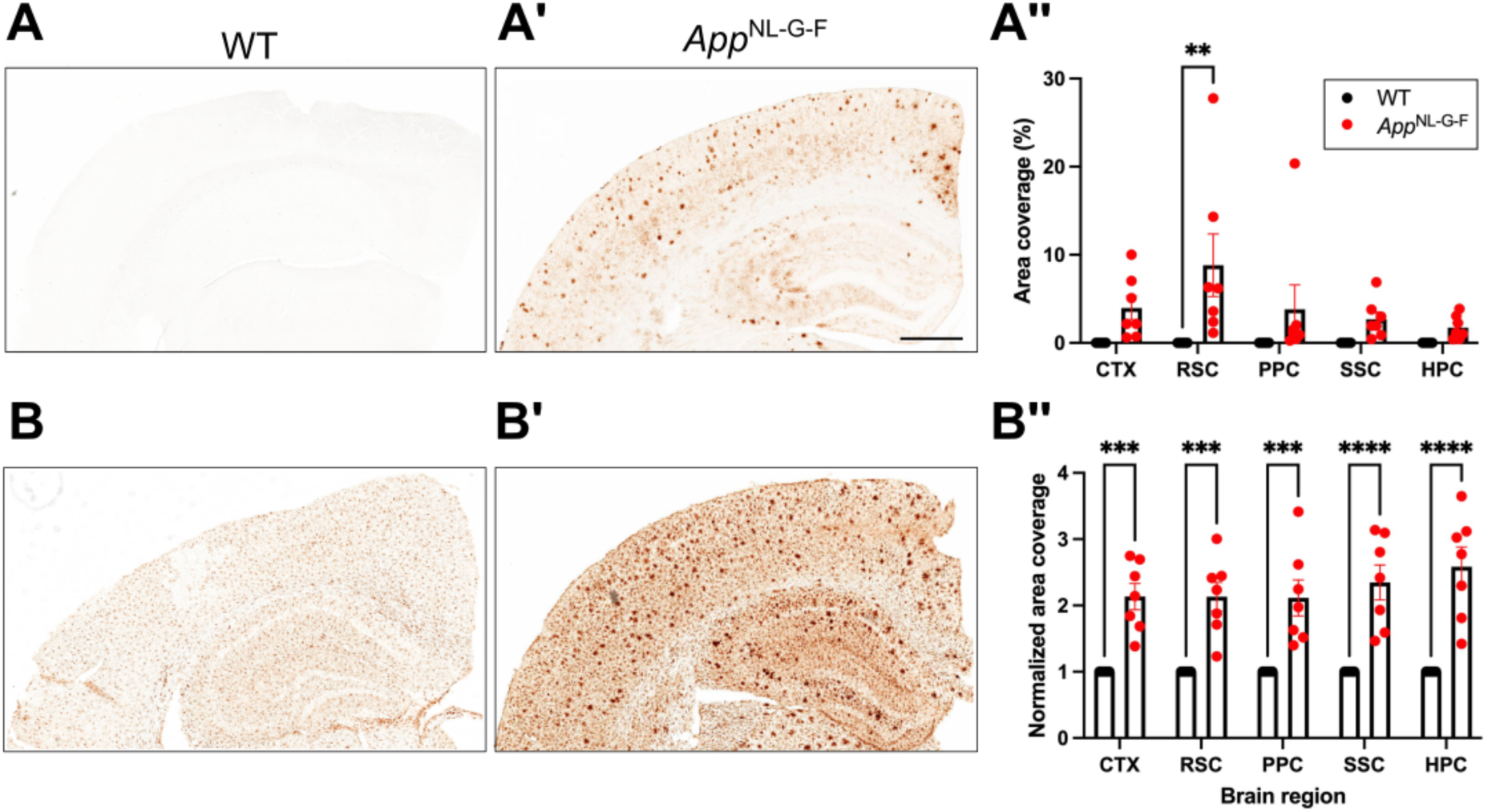
(A) Representative images of cerebral cortex and hippocampus stained for 82E1-positive Aβ deposits in WT and (A’) 22-month-old *App*^NL-G-F^ mice. A” compares the percent area impacted by Aβ deposition in five brain regions between the two strains. (B) Iba1 immunostaining of the cerebral cortex and hippocampus of WT and (B’) 22-month-old *App*^NL-G-F^ mice. (B”) Proportion of 5 brains regions showing severe microgliosis in *App*^NL-G-F^ compared to WT mice. Abbreviations: CTX, cortex; RSC, retrosplenial cortex; PPC, posterior parietal cortex; SSc, somatosensory cortex; HPC, hippocampus. Scale bar: 700 µm. Two-way ANOVA revealed significant effect of genotype for both Aβ and Iba1. \*\**p* < 0.01 and \*\*\**p* < 0.001 based on Šidák’s multiple comparisons test.

### EEG Spectra in App^NL-G-F^ and WT mice

Figure 3A presents a heat map of the EEG spectra (0-25 Hz) across a 24-h period for a representative individual of each sex and genotype. In comparison to age-matched WT littermates, the EEG spectra of the female *App*^NL-G-F^ mouse exhibited less spectral power in the 0-5 Hz range during the dark phase, suggesting less sleep. During the light phase, the sleep bouts of this female also appeared to be shorter in duration than in any of the other three mice in Figure 3A.

**Figure 3.**
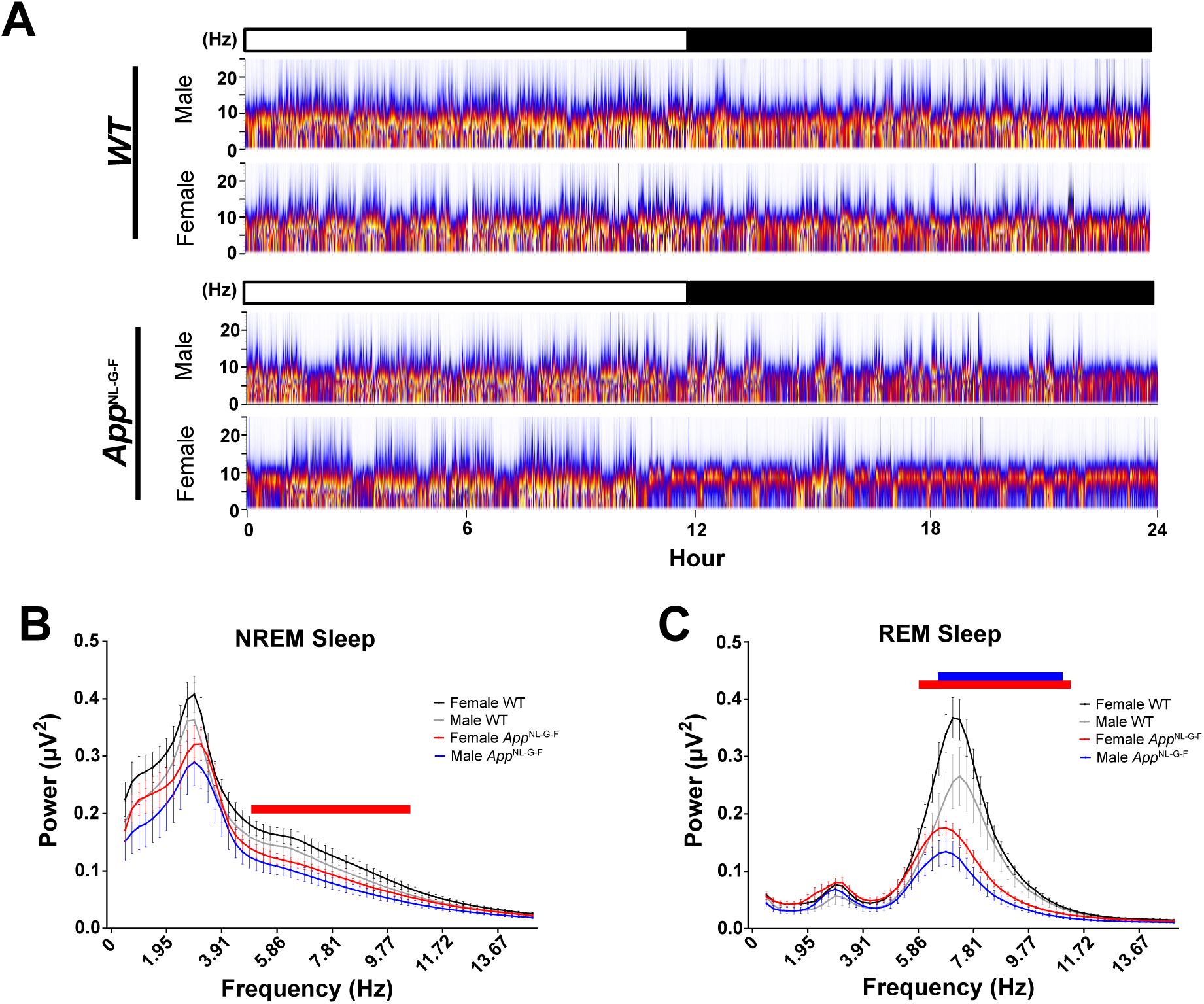
A. EEG spectrograms (0-25 Hz) across the 24-h period for a representative individual of each sex and each genotype. The light and dark phases are indicated in the bar at the top of the spectrogram. Note lower spectral power (indicated by the presence of cooler colors in the 0-5Hz delta range) during the dark phase in the female *App*^NL-G-F^ mouse, suggesting that female mice of this strain sleep less during the dark phase. In contrast, the more consolidated periods with higher power (indicated by the presence of warmer colors in the delta/theta range) in the *App*^NL-G-F/NL-G-F^ males and females during the light period may indicate more consolidated sleep periods during this phase. EEG spectral power (0.5-15 Hz) during (B) NREM and (C) REM sleep for male and female WT and *App*^NL-G-F^ mice at 18.3-24.8 months of age. Red bars indicate the frequency range over which female WT and *App*^NL-G-F^ mice are significantly different; blue bars in B indicate the frequency range over which male WT and *App*^NL-G-F^ mice differ. Bars indicate *p* < 0.05 using unpaired t-test.

### Sex differences in state-specific EEG spectral power in App^NL-G-F^ mice at 18-24 months of age

Spectral analysis of the EEG (0.5-15 Hz) was conducted for all epochs of Wake, NREM and REM sleep. As expected, EEG quality during Wake epochs varied depending upon the magnitude of locomotor activity during any given epoch. Because of the size of the dataset (31 mice x 14 days/mouse x 24 h/day x 60 min/h x 6 epochs/min = 3.75 x 10^6^ 10-s epochs), it was not practical to inspect all Wake epochs to eliminate artifacts due to activity that might distort the spectral analysis during Wake. Consequently, the EEG spectra presented in Figures 3B and **3C** are only for NREM and REM sleep, respectively. Two-way ANOVA revealed significant variation in EEG power across bandwidths during NREM sleep (*F*_(59, 1652)_ = 1.590; *p* = 0.0033). *Post hoc* analysis indicated reduced spectral power in the EEG of female *App*^NL-G-F^ mice relative to female WT mice between 4.88 −10.50 Hz during NREM sleep. A similar analysis of the EEG during REM sleep documented reduced spectral power in the EEG between 6.35 – 11.72 Hz for female *App*^NL-G-F^ mice and between 7.08-11.48 Hz for male *App*^NL-G-F^ mice relative to sex-matched WT mice.

### Distribution of Arousal States, Activity and Body Temperature at 14-18 vs.18-22 months of age

Figure 4 presents the time in each state, activity and subcutaneous body temperature (T_sc_) based on 24-h recordings collected from WT and *App*^NL-G-F^ mice at 14-18 months of age and again at 18-22 months. A mixed-effects model ANOVA with Age, Genotype, and Time of Day as factors indicated significant main effects of Genotype and Time of Day for all five of the parameters measured (**Table 1**). There was a main effect of Age on activity and T_sc_ but not for the amounts of Wake, NREM or REM sleep (**Table 1**). At 14-18 months, *App*^NL-G-F^ mice spent significantly more time awake (16.1%; **Figure 4A’)** and less time in both NREM (−18.2%; **Figure 4B’)** and REM (-14.0%; **Figure 4C’)** sleep across the 24-h period than WT mice. By 18-22 months, *App*^NL-G-F^ mice spent 18.8% more time awake, 19.9% less time in NREM and 18.2% less time in REM sleep. The mixed model ANOVA also revealed significant Age x Genotype interactions for both activity and T_sc_ (**Table 1**). *App*^NL-G-F^ mice were hyperactive relative to WT mice at 18-22 months but not at 14-18 months; Figures 4D and **4D’** document this notable age-related effect on activity which likely affected the distribution of arousal states at 18-22 months. Figure 4E presents the diurnal rhythm of subcutaneous T_sc_ in both genotypes at both ages and **Figure 4E’** shows that the mean T_sc_ across the 24-h period declines with age in both genotypes, as indicated by the main effect of Age in the mixed model ANOVA.

**Figure 4.**
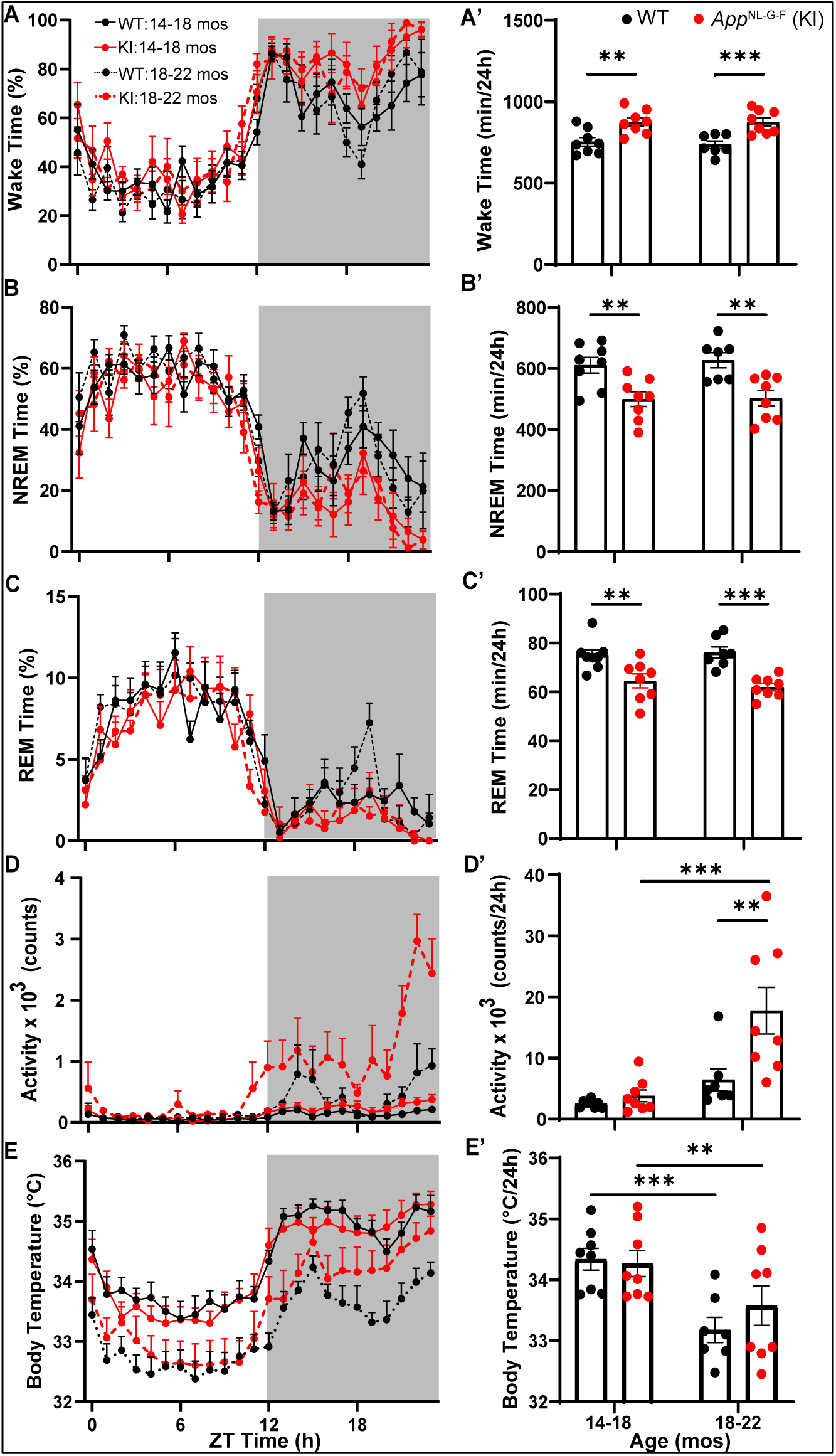
Hourly percent time spent in (A) Wake, (B) NREM sleep and (C) REM sleep as well as activity counts (D) and subcutaneous body temperature (E) across a 24-h recording period in Cohort 1 WT and *App*^NL-G-F^ mice recorded at 14-18 months and at 18-22 months of age. Panels A’-E’ present the corresponding total amounts of each state or physiological parameter for the same 24-h period at each age. \*\**p* < 0.01 and \*\*\**p* < 0.001 based on between group *post hoc* comparisons using Fisher’s LSD test.

**Table 1.** Effects of Age, Genotype, Time of Day and their interactions on vigilance state amounts and activity during baseline for Cohort 1 determined by a mixed-effects model.

|  | Wake |  | NREM |  | REM |  | Activity |  | Body Temperature |  |
| --- | --- | --- | --- | --- | --- | --- | --- | --- | --- | --- |
| Age | F(1, 312) = 0.2362 | NS | F(1, 312) = 0.34 | NS | F(1, 312) = 0.03613 | NS | F(1, 312) = 89.75 | $p < 0.0001$ | F(1, 312) = 515.6 | $p < 0.0001$ |
| Genotype | F(1, 336) = 33.18 | $p < 0.0001$ | F(1, 336) = 35.08 | $p < 0.0001$ | F(1, 336) = 11.14 | $p = 0.0009$ | F(1, 336) = 37.76 | $p < 0.0001$ | F(1, 336) = 5.264 | $p = 0.0224$ |
| Time of Day | F(23, 336) = 31.84 | $p < 0.0001$ | F(23, 336) = 29.83 | $p < 0.0001$ | F(23, 336) = 28.00 | $p < 0.0001$ | F(23, 336) = 7.422 | $p < 0.0001$ | F(23, 336) = 16.49 | $p < 0.0001$ |
| Time x Age | F(23, 312) = 1.062 | NS | F(23, 312) = 1.073 | NS | F(23, 312) = 0.7956 | NS | F(23, 312) = 4.969 | $p < 0.0001$ | F(23, 312) = 0.8463 | NS |
| Time x Genotype | F(23, 336) = 1.036 | NS | F(23, 336) = 1.150 | NS | F(23, 336) = 0.6370 | NS | F(23, 336) = 2.046 | $p = 0.0035$ | F(23, 336) = 0.4083 | NS |
| Age x Genotype | F(1, 312) = 0.2386 | NS | F(1, 312) = 0.1866 | NS | F(1, 312) = 0.3650 | NS | F(1, 312) = 28.06 | $p < 0.0001$ | F(1, 312) = 32.46 | $p < 0.0001$ |
| Time x Age x Genotype | F(23, 312) = 1.090 | NS | F(23, 312) = 1.045 | NS | F(23, 312) = 1.239 | NS | F(23, 312) = 1.722 | $p = 0.0224$ | F(23, 312) = 0.4310 | NS |

Abbreviations: NS, Not significant.

Figure 5 presents the time spent by WT and *App*^NL-G-F^ mice in each state, activity and T_sc_ during the light and dark phases at 14-18 months and 18-22 months of age. As expected, the distribution of arousal states differed between the light and dark phases in both genotypes with more Wake during the dark phase and more NREM and REM sleep in the light phase in this nocturnal species. At 14-18 months, the 16.1% greater levels of Wake and lower levels of NREM and REM sleep in the *App*^NL-G-F^ mice evident in Figure 4 were exclusively due to differences during the dark phase (**Figure 5A’, 5B’ and 5C’).** In contrast, the changes in arousal states at 18-22 months of age in *App*^NL-G-F^ mice relative to WT mice evident in Figure 4 were due to increased amounts of Wake and reduced NREM sleep that occurred in both the light (Figures 5A**, 5B**) and dark phases (**Figures 5A’ and 5B’**). Undoubtedly, a major factor that underlies the increased wake and reduced sleep in *App*^NL-G-F^ mice at 18-22 months of age was increased activity during the dark phase as the mice aged (Figures 4D**, 5D’**). Mixed-effects model ANOVA with Age and Genotype as factors confirmed a main of effect of Age on activity during the light phase (*p* = 0.0359; *F*_(1,13)_ = 5.478) and a stronger effect during the dark phase (*p* = 0.0007; *F*_(1,13)_ = 19.40) as well as both a Genotype and Age x Genotype interaction during the dark phase. Mixed model ANOVA also revealed a significant effect of Age on T_sc_ during both the light (*p* = 0.0003; *F*_(1,13)_ = 24.41) and dark (*p* < 0.0001; *F*_(1,13)_ = 37.95) phases; mean T_sc_ declined with age in both genotypes during both the light (Figure 5E) and dark (**Figure 5E’**) phases.

**Figure 5.**
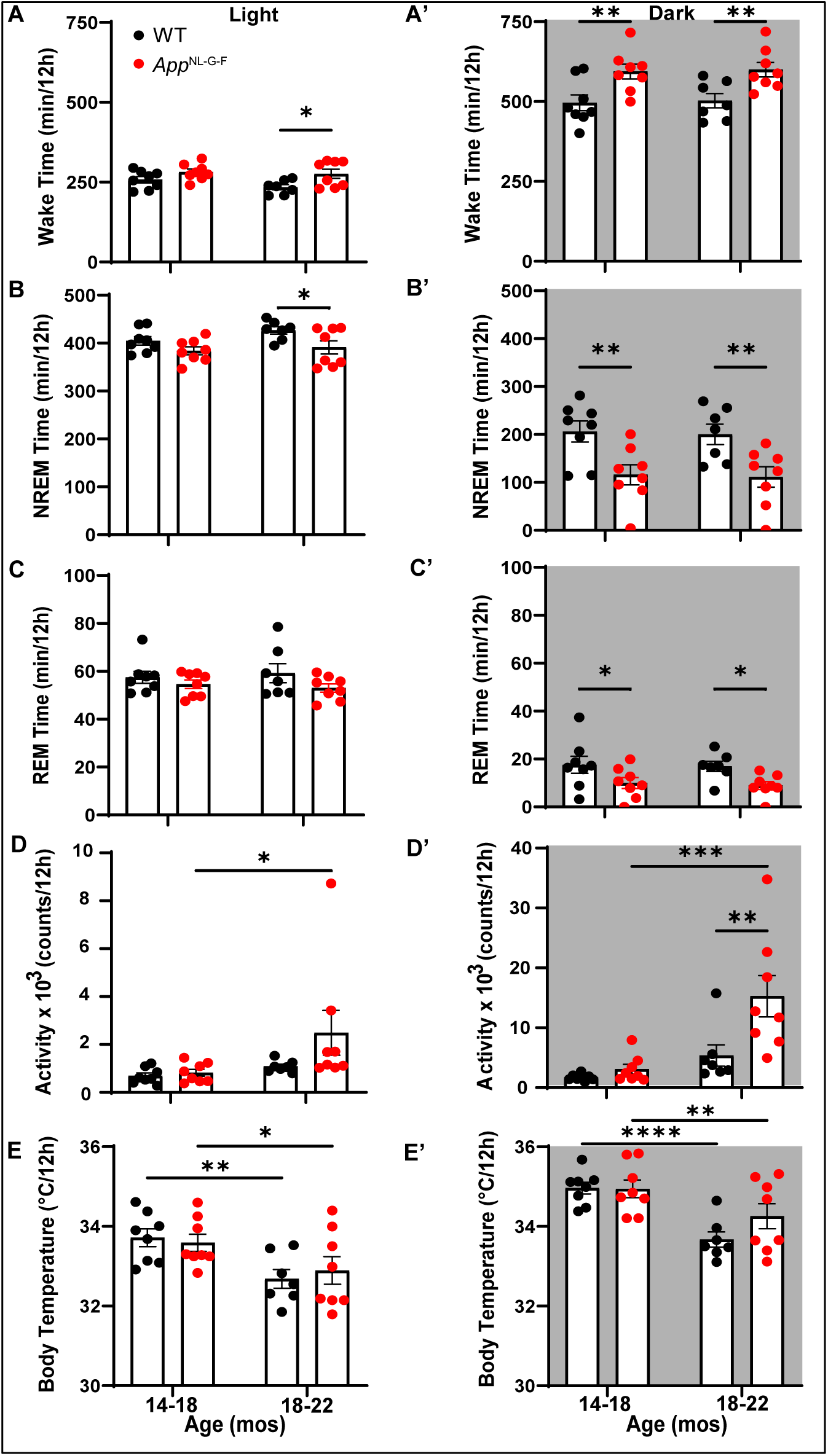
Amount of time in (A) Wake, (B) NREM sleep and (C) REM sleep as well as total activity counts (D) and subcutaneous body temperature (E) during the 12-h light phase in WT and *App*^NL-G-F^ mice at 14-18 months and at 18-22 months of age. Panels A’-E’ present the corresponding results for the 12-h dark phase at each age. \**p* < 0.05, \*\**p* < 0.01, \*\*\**p* < 0.001 and \*\*\*\**p* < 0.0001 based on between group *post hoc* comparisons using Fisher’s LSD test.

### Sleep Architecture in Cohort 1 mice at 14-18 vs. 18-22 months of age

The EEG/EMG data summarized in Figure 4 were analyzed for conventional measures of sleep architecture, i.e., the number of bouts/state and mean bout duration/state. A mixed-effects model ANOVA with Age, Genotype and Time of Day as factors indicated significant main effects of Genotype and Time of Day on the number of Wake, NREM and REM sleep bouts (*p* < 0.0001 for all states; **Table 2**) but, as in Table 1, there was no significant effect of Age on the number of bouts for any of these states.

**Table 2.** Effects of Age, Genotype, Age, Time of Day and their interactions on the number of bouts of each state during baseline for Cohort 1 determined by a mixed-effects model.

|  | Wake |  | NREM |  | REM |  |
| --- | --- | --- | --- | --- | --- | --- |
| Age | F(1, 312) = 1.122 | NS | F(1, 312) = 0.4082 | NS | F(1, 312) = 0.3319 | NS |
| Genotype | F(1, 336) = 53.45 | $p < 0.0001$ | F(1, 336) = 53.17 | $p < 0.0001$ | F(1, 336) = 41.92 | $p < 0.0001$ |
| Time of Day | F(23, 336) = 6.050 | $p < 0.0001$ | F(23, 336) = 8.087 | $p < 0.0001$ | F(23, 336) = 21.38 | $p < 0.0001$ |
| Time x Age | F(23, 312) = 0.7334 | NS | F(23, 312) = 0.8242 | NS | F(23, 312) = 0.7154 | NS |
| Time x Genotype | F(23, 336) = 0.8157 | NS | F(23, 336) = 0.8051 | NS | F(23, 336) = 0.8121 | NS |
| Age x Genotype | F(1, 312) = 0.07494 | NS | F(1, 312) = 0.004878 | NS | F(1, 312) = 0.1341 | NS |
| Time x Age x Genotype | F(23, 312) = 0.9135 | NS | F(23, 312) = 0.9417 | NS | F(23, 312) = 0.7453 | NS |

Abbreviations: NS, Not significant.

On the other hand, mixed-effects model ANOVA on the duration of Wake, NREM and REM sleep bouts with Age, Genotype and Time of Day yielded more complex results. Although there was a main effect of Time of Day for all 3 states (*p* < 0.0001; **Table 3),** Genotype was significant only for Wake bout duration (*p* < 0.0001; **Figure 6A’**) but there were Genotype x Time of Day interactions for all 3 states (Wake: *p* < 0.0001; NREM: *p* = 0.0015; and REM: *p* = 0.0054). There were also main effects of Age for both NREM (*p* = 0.0294) and REM (*p* = 0.0337) bout duration (**Table 3)**.

**Figure 6.**
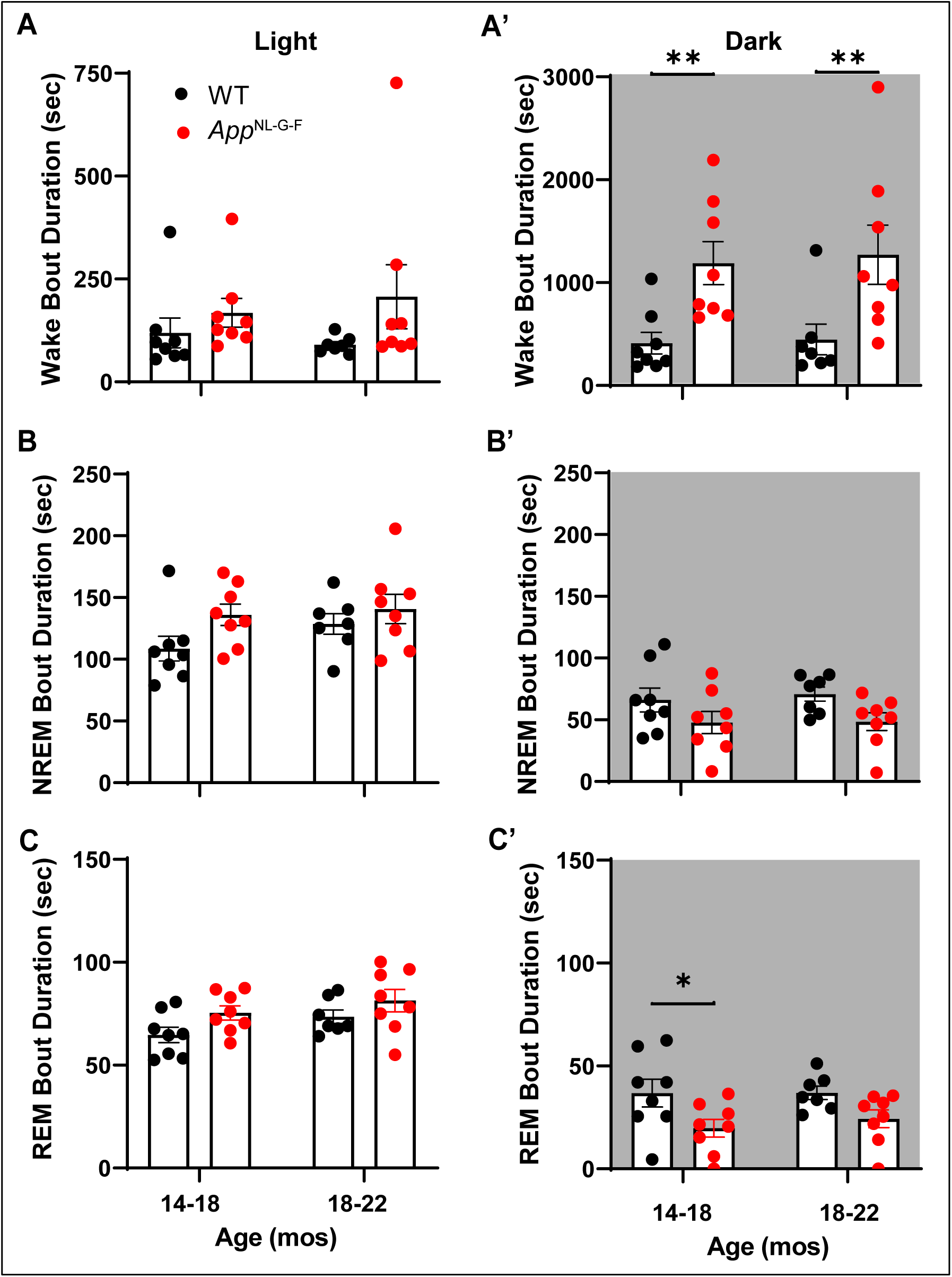
Mean bout duration during (A) Wake, (B) NREM and (C) REM sleep during the 12-h light phase in WT and *App*^NL-G-F^ mice at 14-18 months and at 18-22 months of age. Panels A’-C’ present the corresponding results for the 12-h dark phase at each age. \**p* < 0.05 and \*\**p* < 0.01 based on between group *post hoc* comparisons using Fisher’s LSD test.

**Table 3.** Effects of Age, Genotype, Time of Day and their interactions on bout durations during baseline for Cohort 1 mice determined by a mixed-effects model.

|  | Wake |  | NREM |  | REM |  |
| --- | --- | --- | --- | --- | --- | --- |
| Age | F(1, 312) = 0.3363 | NS | F(1, 312) = 4.791 | $p = 0.0294$ | F(1, 312) = 4.548 | $p = 0.0337$ |
| Genotype | F(1, 336) = 50.67 | $p < 0.0001$ | F(1, 336) = 0.0058 | NS | F(1, 336) = 1.311 | NS |
| Time of Day | F(23, 336) = 9.544 | $p < 0.0001$ | F(23, 336) = 19.50 | $p < 0.0001$ | F(23, 336) = 18.45 | $p < 0.0001$ |
| Time x Age | F(23, 312) = 0.7740 | NS | F(23, 312) = 0.9393 | NS | F(23, 312) = 0.7694 | NS |
| Time x Genotype | F(23, 336) = 2.876 | $p < 0.0001$ | F(23, 336) = 2.191 | $p = 0.0015$ | F(23, 336) = 1.975 | $p = 0.0054$ |
| Age x Genotype | F(1, 312) = 0.3289 | NS | F(1, 312) = 1.988 | NS | F(1, 312) = 0.02686 | NS |
| Time x Age x Genotype | F(23, 312) = 0.3204 | NS | F(23, 312) = 0.5389 | NS | F(23, 312) = 0.6711 | NS |

Abbreviations: NS, Not significant.

Figure 6 presents the mean Bout Duration for each state during the light and dark phases. Although the mean Bout Duration did not differ between genotypes for any state during the light phase (Figures 6A**, B, and C**), the mean Wake Bout Duration during the dark phase was more than twice as long for the *App*^NL-G-F^ mice at both ages (14-18 months: *p* = 0.0099; 18-22 months: *p* = 0.0064; **Figure 6A’)** and the mean REM Bout Duration for *App*^NL-G-F^ mice was 46.3% shorter at 14-18 months of age (*p* = 0.0195; **Figure 6C’)**. With one exception, the distribution of mean Wake Bout durations of *App*^NL-G-F^ mice during the dark phase at 18-22 months of age did not overlap with that of WT mice of the same age (**Figure 6A’)**, which led us to investigate the distribution of Wake Bout Durations in more detail. The entire 14-day recording of 18-22 month old WT (n=7) and *App*^NL-G-F^ (n=8) mice in Cohort 1 was scored using Somnivore and the number Wake Bouts were binned in durations of <60 sec, 60-100 sec, 100-140 sec, 140-180 sec, 180-220 sec, 220-260 sec and >260 sec. As indicated in Figure 7A, *App*^NL-G-F^ mice had significantly fewer short (<60 sec) Wake Bouts than WT mice (*p* = 2.68 x 10^-6^) but more longer (>260 sec) Wake Bouts (*p* = 1.31 x 10^-6^). These longer Wake Bouts are likely due to the increased activity of *App*^NL-G-F^ mice during the dark phase at 18-22 months (Figures 4D **and 5D’**). Figures 7B and **7C** show that, although activity increases for individuals of both genotypes with age, the age-dependent activity increase was greater in *App*^NL-G-F^ mice during both the light and dark phases.

**Figure 7.**
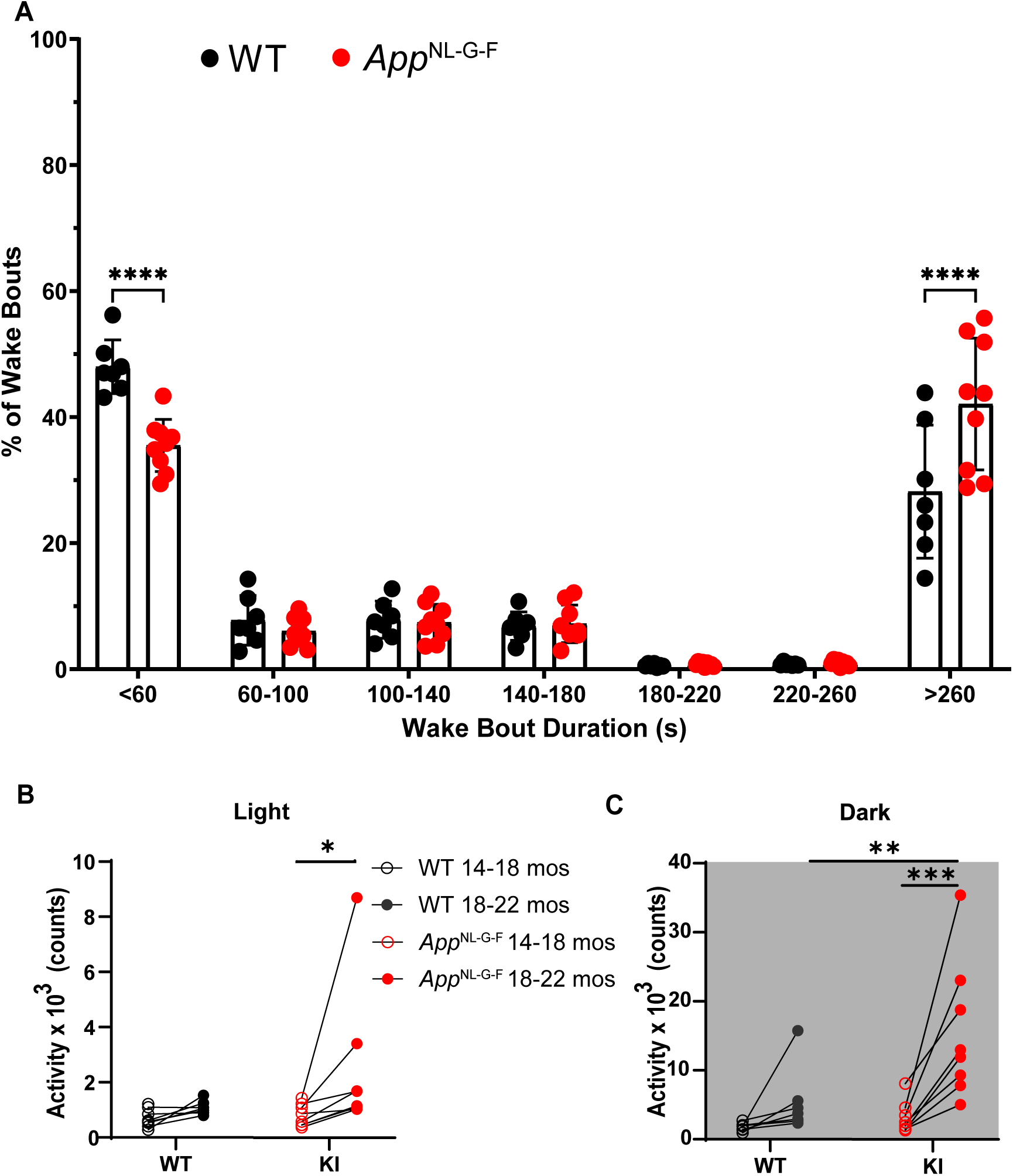
(A) Distribution of Wake Bouts binned in durations varying from <60 sec to >260 sec in 18-22 month old WT and *App*^NL-G-F^ mice. (B) Activity counts per 24-h period in WT and *App*^NL-G-F^ mice during the light phase at both ages. Lines connect the activity of levels of each individual studied at the two ages. (C) Activity counts per 24-h period in WT and *App*^NL-G-F^ mice during the dark phase at both ages. \**p* < 0.05, \*\**p* < 0.01, \*\*\**p* < 0.001 and \*\*\*\**p* < 0.0001 based on between group *post hoc* comparisons using Fisher’s LSD test.

### Multiple Sleep Latency Test and Response to 6-h Sleep Deprivation

The results presented above indicated that *App*^NL-G-F^ mice had less sleep than WT littermates, particularly in the dark phase when Wake bouts were of longer duration, and that activity increased with age. To determine whether the level of sleepiness differed in the two strains, Cohort 1 mice underwent a murine MSLT at 18-22 months of age (Figure 1). **Figure S1** shows that there were no significant differences between the two strains in the latency to or accumulation of NREM or REM sleep during the five 20 min nap opportunities. Consequently, *App*^NL-G-F^ mice at 18-22 months of age appear to be no sleepier than age-matched WT littermates despite having less sleep and greater activity per day, particularly during the dark phase.

To assess the integrity of the sleep homeostat, mice of both strains also underwent a 6-h sleep deprivation (SD) followed by a recovery sleep (RS) opportunity at 18-22 months of age.

Figure 8 shows that, after cessation of the 6-h SD, NREM time increased in *App*^NL-G-F^ mice for several hours during the recovery phase (Figure 8B). Although NREM time did not increase during any particular RS hour in WT mice (Figure 8A), the cumulative amount of NREM sleep over the first 5-h of the RS phase was significantly increased in both genotypes relative to the baseline recorded 24-h earlier (**Figure S2B and S2B’).** REM sleep amounts did not change during the 5-h RS period during the light phase in either genotype (Figures 8C**, 8D and Figure S2C)**. During the subsequent dark phase, however, REM sleep amounts were greater in both *App*^NL-G-F^ and WT mice compared to baseline (**Figure S2C’)**, indicating a delayed recovery of REM sleep. Both strains showed the expected increase in NREM delta power during the first hour after SD cessation that gradually declined over the subsequent 4-h (Figures 8E**, 8F**). Thus, the homeostatic response to SD appears to be intact in *App*^NL-G-F^ mice even at 18-22 months of age.

**Figure 8.**
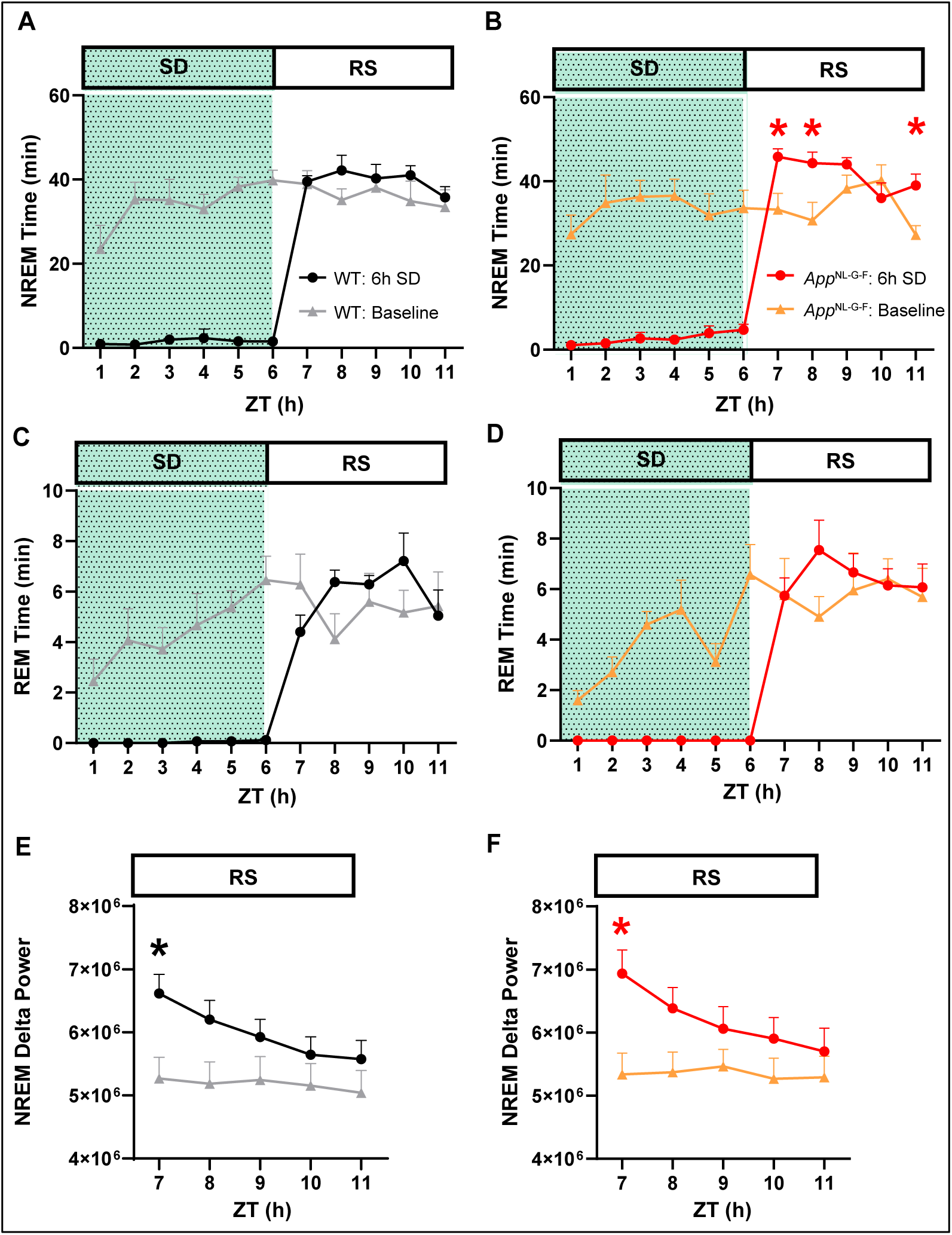
Comparison of homeostatic response to 6-h sleep deprivation in 18-22 month old WT and *App*^NL-G-F^ mice. (A) Hourly amounts of NREM sleep from ZT0 to ZT12 in *App*^WT/WT^ mice on a baseline day (gray line) vs. a day on which 6-h sleep deprivation (SD) occurred (black line) from ZT0-6 (shaded area) followed by a 5-h recovery sleep (RS) opportunity from ZT6-11. (B) Same as A but for *App*^NL-G-F^ mice. (C, D) Same as A and B but for REM sleep. (E, F) EEG delta power in NREM sleep during the RS period (ZT6-11) after the 6-h SD (ZT0-6) compared to the baseline day in (E) WT and (F) *App*^NL-G-F^ mice. Values are mean ± SEM. * *p* < 0.05 based on between condition *post hoc* comparisons using Fisher’s LSD test.

### Comparison of state-specific EEG spectra in Cohort 1 mice at 14-18 vs. 18-22 months of age

Figure 9 compares the EEG Spectral Power Density during Wake, NREM and REM sleep for WT vs. *App*^NL-G-F^ mice at 14-18 vs.18-22 months of age. Mixed-model ANOVA revealed a main effect of Age for all bandwidths during Wake except high gamma. Spectral power density during Wake declined with age in *App*^NL-G-F^ mice for 4 of the 5 bandwidths and for WT mice in 3 of the 5 bandwidths (Figures 9D-G), the exception being the delta range which increased with age for both genotypes (Figure 9C). Spectral power density in the alpha range during Wake was significantly lower in the *App*^NL-G-F^ mice at 18-22 months compared to WT (Figure 9E). The gamma/delta ratio decreased with age in both genotypes (Figure 9H).

**Figure 9.**
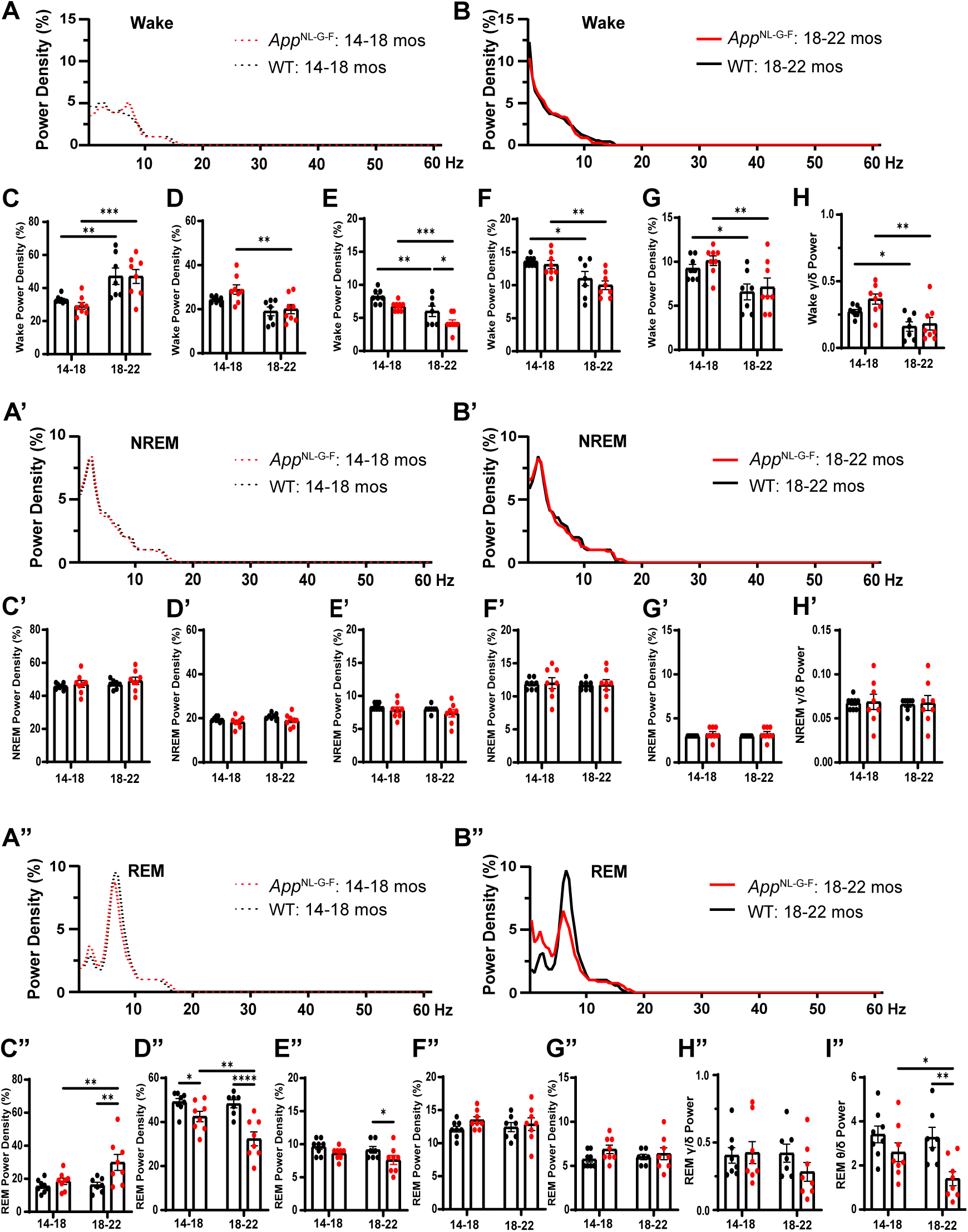
EEG power density during Wake, NREM and REM sleep in WT (black) and *App*^NL-G-F^ (red) mice at 14-18 months (A, A’ and A”) and at 18-22 months (B, B’ and B”). Panels C-H present the EEG Power Density during Wake at both ages in the delta (C), theta (D), alpha (E), beta (F), low gamma (G) ranges and the gamma/delta ratio (H). Panels C’-H’ provide comparable information for NREM sleep. Panels C’’-H’’ provide similar information for REM sleep as well as the theta/delta ratio (I”). * *p* < 0.05, ** *p* < 0.01, *** *p* < 0.001 and **** *p* < 0.0001 based on Fisher’s LSD test after mixed-effects ANOVA.

The spectral power density during NREM sleep was unchanged with age in both genotypes (**Figure 9C’-H’**).

During REM sleep, however, there were significant effects of both Age and Genotype for the delta (**Figure 9C’’**) and theta (**Figure 9D’’**) bands as well as a Genotype effect for the alpha band in which *App*^NL-G-F^ mice exhibited lower power density than WT mice at 18-22 months (**Figure 9E’’**). Spectral power density in the delta range during REM sleep was greater in *App*^NL-G-F^ compared to WT mice at 18-22 months (**Figure 9C’’**) but reduced in the theta band for *App*^NL-G-F^ mice at both ages (**Figure 9D’’**) as also occurs in younger *App*^NL-G-F^ mice (Calafate et al., 2023; Maezono et al., 2020). Moreover, the theta/delta ratio declined with age in *App*^NL-G-F^ mice and was reduced in 18-22 month old *App*^NL-G-F^ *vs.* WT mice (Figure 9I**”).**

*Sex Differences in State Amounts in App*^NL-G-F^ *vs. WT mice at 18-24 months of age*

To undertake a rigorous statistical comparison of *App*^NL-G-F^ and WT mice at an advanced age, the EEG/EMG data collected from the recording of Cohort 1 mice at 18-22 months of age was combined with the recordings collected from Cohort 2 mice. This combination resulted in 31 mice across 4 groups: a female *App*^NL-G-F^ group (N=8) ranging from 20.6-23.8 mos (22.0±0.4 mos), a female WT group (N=6) ranging from 19.8-24.6 mos (22.1±0.6 mos), a male *App*^NL-G-F^ group (N=9) ranging from 18.3-24.8 mos (21.1±0.8 mos), and a male WT group (N=8) ranging from 18.3-24.8 mos (21.4±1.0 mos).

Figure 10 shows the average amount of Wake (A), NREM (B), and REM (C) sleep on 14 consecutive days for the 31 mice recorded when they ranged from 18.3-24.8 months of age. *App*^NL-G-F^ mice spent more time awake and less time in both NREM and REM sleep compared to the WT mice. Although a long wake/short sleep phenotype was consistent between *App*^NL-G-F^ males and females, it was more apparent in females.

**Figure 10.**
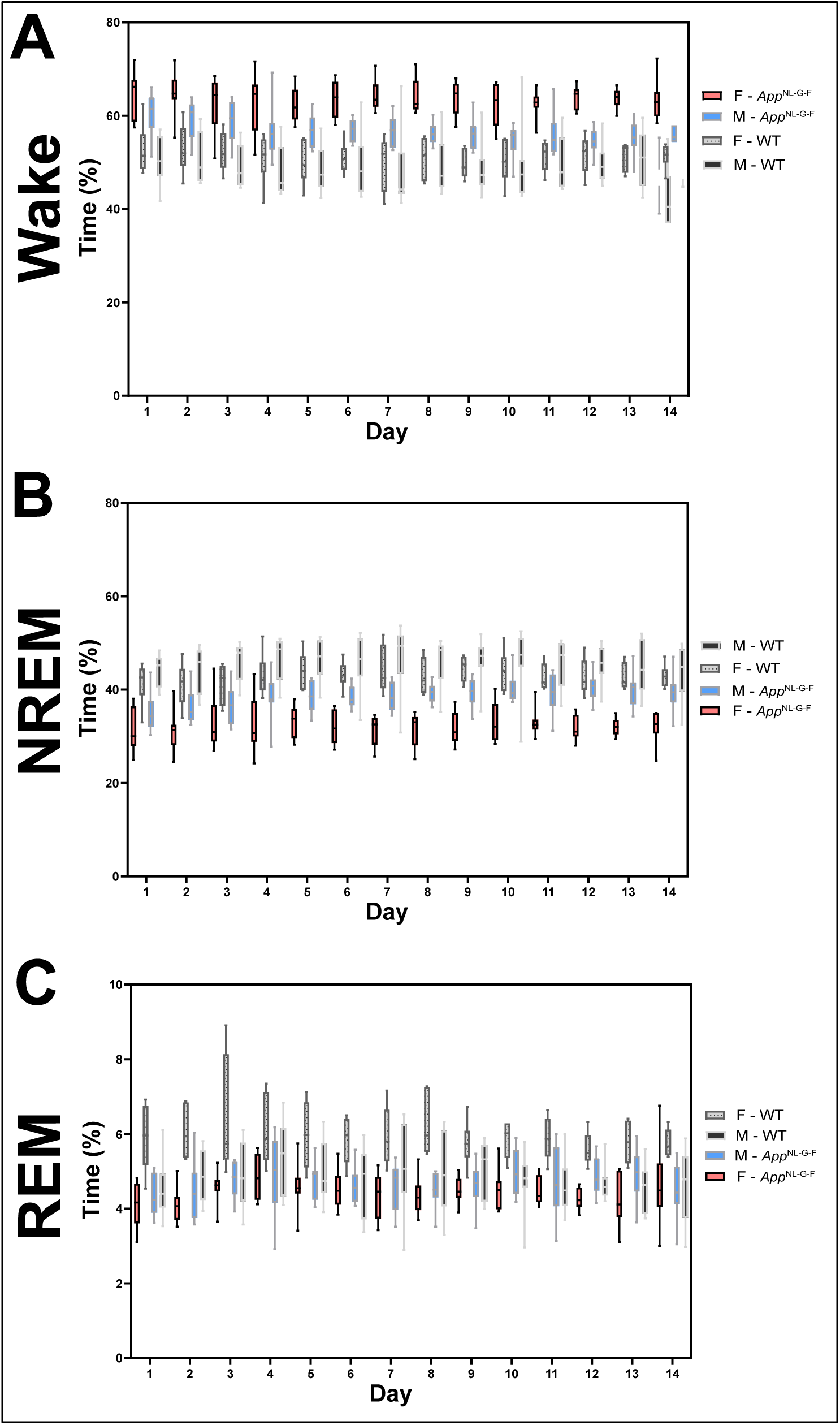
Mean percentage of time spent in Wake (A), NREM (B), and REM (C) sleep on 14 consecutive days for the 31 mice recorded when they ranged from 18.3-24.8 months of age.

Figure 11 presents the amount of each state for both strains separated by sex across the entire 24-h period as well as for the light and dark phases. Two-way ANOVA revealed significant effects of Genotype and Sex for the amounts of Wake, NREM, and REM sleep across the entire 24-h period (**Table 4**); however, a significant Genotype x Sex interaction was only indicated for the amount of REM sleep (*F*_(1, 27)_ = 10.95; *p* = 0.0027). *Post hoc* tests revealed that both male and female *App*^NL-G-F^ mice have more Wake (Fig. 11A) and less NREM sleep (Fig. 11**A****’**) than WT littermates; however, only female *App*^NL-G-F^ mice have less REM sleep (Fig. 11**A****’’**). These genotype and sex effects were also evident during the dark phase (**Figs. 11C, C’ and C”)** but, during the light phase, were only present in the *App*^NL-G-F^ females who continued to exhibit an insomnia-like phenotype with more Wake (Fig. 11B) and less NREM (Fig. 11**B****’**) and REM (Fig. 11**B****’’**) sleep. Overall, female *App*^NL-G-F^ mice were more severely affected than males.

**Figure 11.**
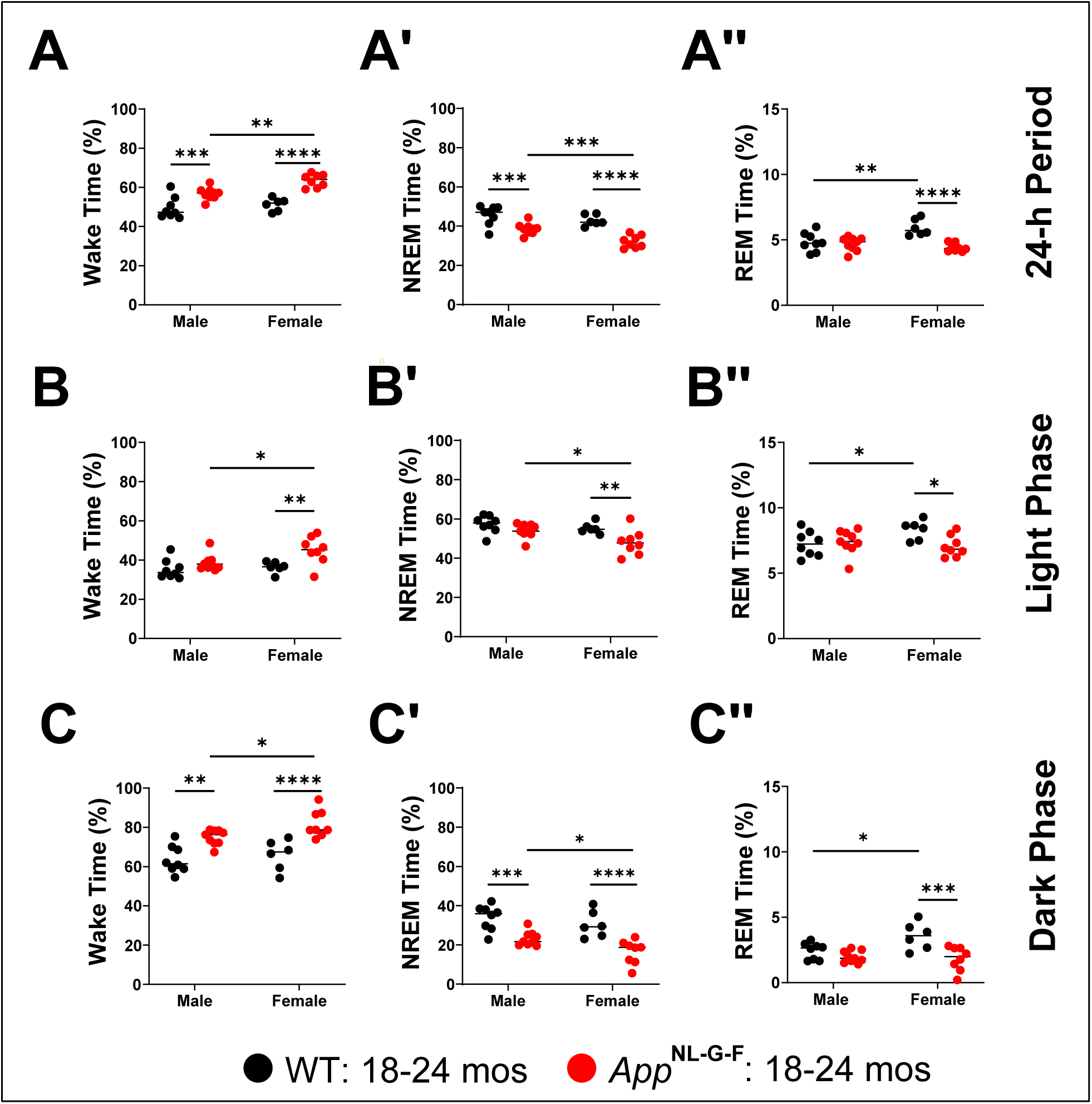
Percent time in sleep/wake states for male and female WT and *App*^NL-G-F^ mice at 18.3-24.8 months of age. Percent time for Wake (A-C), NREM (A’-C’), and REM sleep (A’’-C’’) averaged across the 14 24-h periods recorded (A-A’’) and for the corresponding light (B-B’’) and dark phases (C-C’’). Across the 24-h periods, *App*^NL-G-F^ mice of both sexes exhibited a greater amount of Wake (A) and decreased amounts of REM sleep (A’’) in comparison to their age-matched WT littermate controls. While this effect is most pronounced in the dark phase in both sexes (C-C’’), it is also evident during the light phase in female mice (B-B’’). \**p* < 0.05, \*\**p* < 0.01, \*\*\**p* < 0.001 and \*\*\*\**p* < 0.0001 based on between group *post hoc* comparisons using Fisher’s LSD test.

**Table 4.**
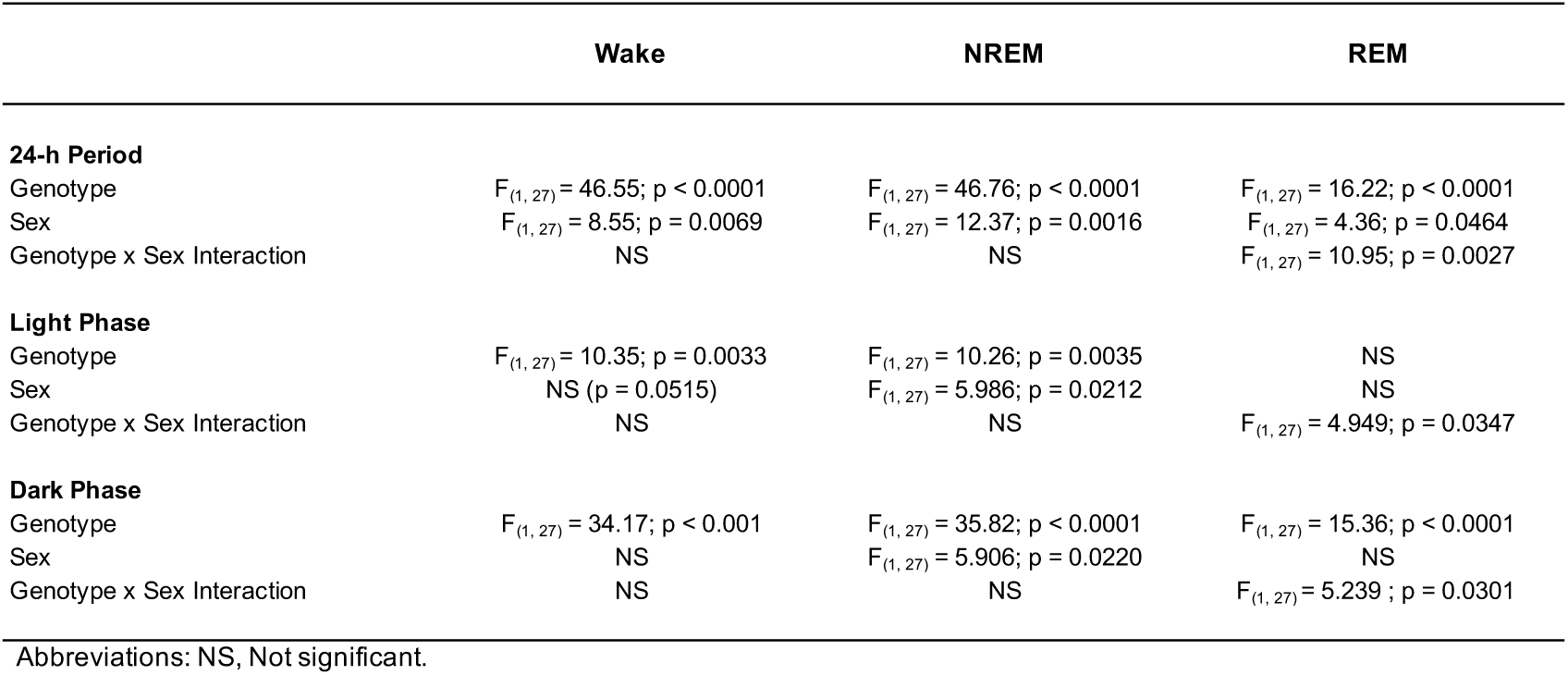
Effects of Genotype, Sex, and Genotype x Sex interaction on vigilance state amounts during baseline and during the light and dark phases.

|  | Wake | NREM | REM |
| --- | --- | --- | --- |
| **24-h Period** |  |  |  |
| Genotype | $F_{(1, 27)} = 46.55; p < 0.0001$ | $F_{(1, 27)} = 46.76; p < 0.0001$ | $F_{(1, 27)} = 16.22; p < 0.0001$ |
| Sex | $F_{(1, 27)} = 8.55; p = 0.0069$ | $F_{(1, 27)} = 12.37; p = 0.0016$ | $F_{(1, 27)} = 4.36; p = 0.0464$ |
| Genotype x Sex Interaction | NS | NS | $F_{(1, 27)} = 10.95; p = 0.0027$ |
| **Light Phase** |  |  |  |
| Genotype | $F_{(1, 27)} = 10.35; p = 0.0033$ | $F_{(1, 27)} = 10.26; p = 0.0035$ | NS |
| Sex | NS ($p = 0.0515$) | $F_{(1, 27)} = 5.986; p = 0.0212$ | NS |
| Genotype x Sex Interaction | NS | NS | $F_{(1, 27)} = 4.949; p = 0.0347$ |
| **Dark Phase** |  |  |  |
| Genotype | $F_{(1, 27)} = 34.17; p < 0.001$ | $F_{(1, 27)} = 35.82; p < 0.0001$ | $F_{(1, 27)} = 15.36; p < 0.0001$ |
| Sex | NS | $F_{(1, 27)} = 5.906; p = 0.0220$ | NS |
| Genotype x Sex Interaction | NS | NS | $F_{(1, 27)} = 5.239; p = 0.0301$ |

Abbreviations: NS, Not significant.

### Sex Differences in Bout Durations in App^NL-G-F^ mice at 18-24 months of age

Across the entire 24-h period, there were significant effects of Genotype on the durations of both Wake (*F*_(1, 27)_ = 34.34; *p* < 0.0001) and REM sleep (*F*_(1, 27)_ = 21.89; *p* < 0.0001) bouts, for Sex on Wake Bout Duration (*F*_(1, 27)_ = 10.13; *p* = 0.0037), and a significant Genotype x Sex interaction was only indicated for NREM sleep Bout Duration (*F*_(1, 27)_ = 7.772; *p* = 0.0096; **Table 5**). Wake (Fig. 12A) and REM sleep (Fig. 12**A****’’**) bouts were longer in both sexes of *App*^NL-G-F^ mice than in their WT littermates; NREM bouts were longer only in *App*^NL-G-F^ males (Fig. 12**A****’**).

**Figure 12.**
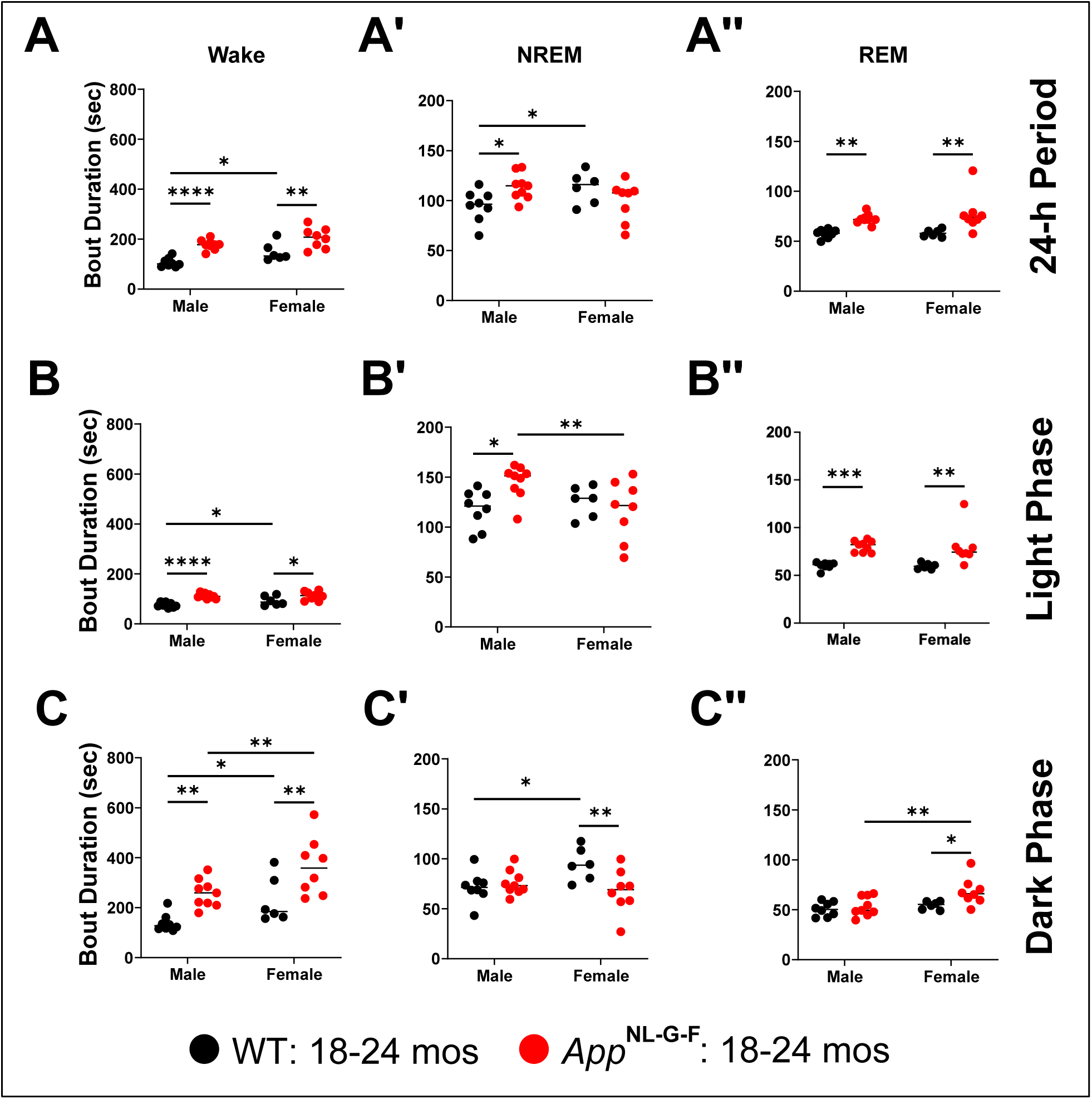
Mean bout durations of sleep/wake states for male and female WT and *App*^NL-G-F^ mice at 18.3-24.8 months of age. Wake (A-C), NREM (A’-C’), and REM sleep (A’’-C’’) bout durations were averaged for the 14 24-h periods recorded (A-A’’) and for the corresponding light (B-B’’) and dark phases (C-C’’). *App*^NL-G-F^ mice exhibit longer Wake (A), NREM (A’) and REM sleep (A’’) bouts in comparison to age-matched WT littermate controls. This effect is largely consistent across both the light (B-B’’) and dark (C-C’’) phases. \**p* < 0.05, \*\**p* < 0.01, \*\*\**p* < 0.001 and \*\*\*\**p* < 0.0001 based on between group *post hoc* comparisons using Fisher’s LSD test.

**Table 5.**
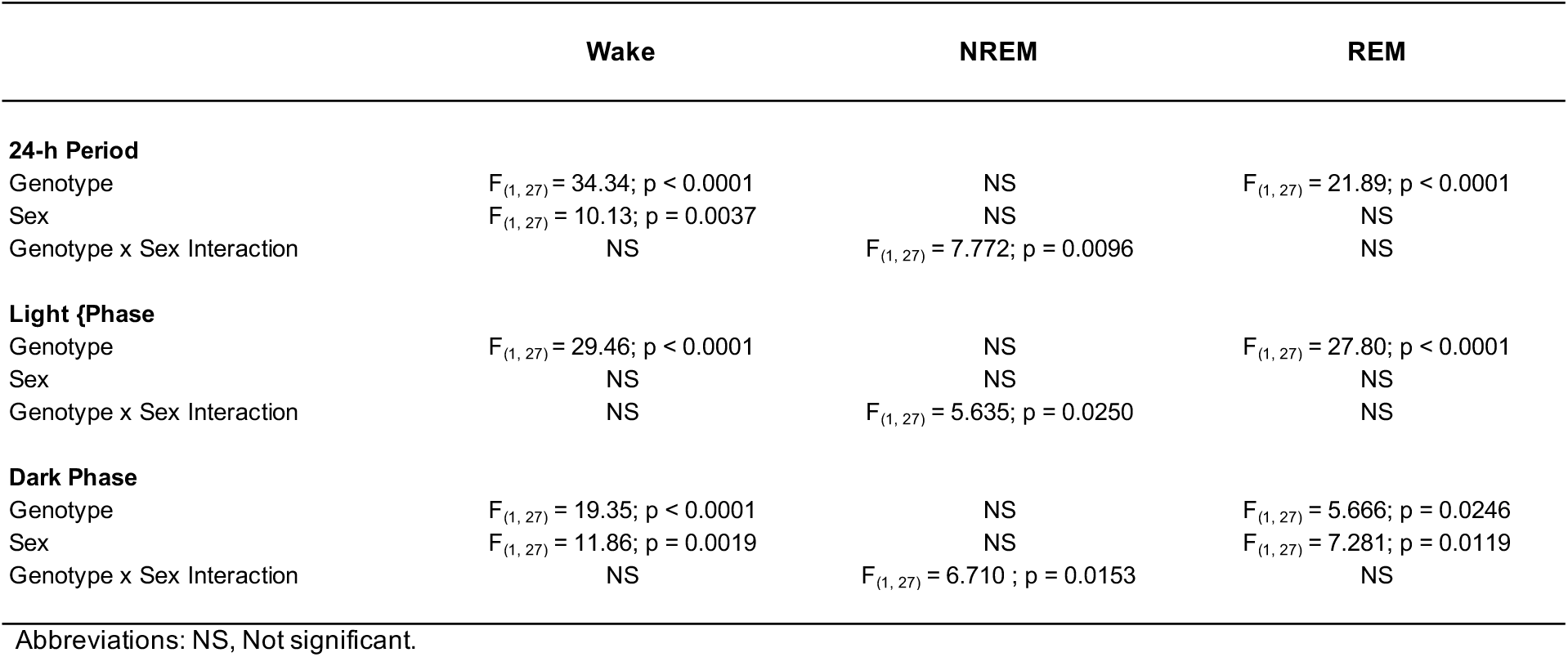
Effects of Genotype, Sex, and Genotype x Sex interaction on bout duration of each vigilance state during baseline and the light and dark phases.

The same patterns were evident during the light phase (**Table 5**): there were significant Genotype effects for Wake (*F*_(1, 27)_ = 29.46; *p* < 0.0001) and REM sleep (*F*_(1, 27)_ = 27.80; *p* < 0.0001) bout durations and a Genotype x Sex interaction for NREM Bout Duration (*F*_(1, 27)_ = 5.635; *p* < 0.0250). Wake (Fig. 12B) and REM sleep (Fig. 12**B****’’**) bouts were longer during the light phase in *App*^NL-G-F^ mice in both sexes than in their WT littermates but NREM bouts were longer only in *App*^NL-G-F^ males (Fig. 12**B****’**).

During the dark phase, there were again significant Genotype effects for Wake (*F*_(1, 27)_ = 19.35; *p* < 0.0001) and REM sleep (*F*_(1, 27)_ = 5.666; *p* < 0.0246) Bout Durations, as well as Sex effects on Wake (*F*_(1, 27)_ = 11.86; *p* = 0.0019) and REM sleep (*F*_(1, 27)_ = 7.281; *p* < 0.0119). Bout Durations, and a Genotype x Sex interaction for NREM Bout Duration (*F*_(1, 27)_ = 6.71; *p* < 0.0153; **Table 5**). As in the light phase, Wake Bout Durations during the dark phase were longer in both sexes of *App*^NL-G-F^ mice than in their WT littermates (*p* < 0.01; Fig. 12C); this was also the case for REM Bout Durations in female *App*^NL-G-F^ mice (*p* < 0.05; Fig. 12C**”**) but not for males. Consistent with an insomnia-like phenotype, NREM Bout Durations were shorter in female *App*^NL-G-F^ mice than in their female WT littermates (*p* < 0.01; Fig. 12**C****’).**

### Sex Differences in the Number of Sleep/Wake Bouts in App^NL-G-F^ mice at 18-24 months of age

Two-way ANOVA revealed significant effects of Genotype for the number of Wake (*F*_(1, 27)_ = 18.34; *p* < 0.0001), NREM (*F*_(1, 27)_ = 21.83; *p* < 0.0001), and REM sleep (*F*_(1, 27)_ = 46.10; *p* < 0.0001) bouts across the entire 24-h period, as well as during the light and dark phases (**Table 6**). There was also a significant main effect of Sex for the number of bouts of Wake (*F*_(1, 27)_ = 7.237; *p* = 0.0121) and NREM sleep (*F*_(1, 27)_ = 6.338; *p* = 0.0181) across the 24-h period and during the dark phase (Wake: *p* < 0.0001; NREM: *p* < 0.0001; **Table 6**). Genotype x Sex interaction effects were significant for number of bouts of Wake (*F*_(1, 27)_ = 6.97; *p* = 0.0136), NREM (*F*_(1, 27)_ = 5.00; *p* = 0.0338), and REM (*F*_(1, 27)_ = 6.79; *p* = 0.0148) sleep across the 24-h period, for Wake (*F*_(1, 27)_ = 6.862; *p* = 0.0143) and NREM sleep (*F*_(1, 27)_ = 4.876; *p* = 0.0359) during the light phase, and for REM (*F*_(1, 27)_ = 5.47; *p* = 0.0269) sleep during the dark phase (**Table 6**).

**Table 6.** Effects of Genotype, Sex, and Genotype x Sex interaction on the number of bouts of each vigilance state during baseline and the light and dark phases.

|  | Wake | NREM | REM |
| --- | --- | --- | --- |
| **24-h Period** |  |  |  |
| Genotype | $F_{(1, 27)} = 18.35; p < 0.0001$ | $F_{(1, 27)} = 21.83; p < 0.0001$ | $F_{(1, 27)} = 46.10; p < 0.0001$ |
| Sex | $F_{(1, 27)} = 7.237; p = 0.0121$ | $F_{(1, 27)} = 6.338; p = 0.0181$ | NS |
| Genotype x Sex Interaction | $F_{(1, 27)} = 6.966; p = 0.0136$ | $F_{(1, 27)} = 4.998; p = 0.0338$ | $F_{(1, 27)} = 6.785; p = 0.0148$ |
| **Light Phase** |  |  |  |
| Genotype | $F_{(1, 27)} = 4.480; p = 0.0437$ | $F_{(1, 27)} = 6.330; p = 0.0181$ | $F_{(1, 27)} = 57.34; p < 0.0001$ |
| Sex | NS | NS | NS |
| Genotype x Sex Interaction | $F_{(1, 27)} = 6.862; p = 0.0143$ | $F_{(1, 27)} = 4.876; p = 0.0359$ | NS |
| **Dark Phase** |  |  |  |
| Genotype | $F_{(1, 27)} = 22.13; p < 0.0001$ | $F_{(1, 27)} = 23.75; p < 0.0001$ | $F_{(1, 27)} = 16.97; p < 0.0001$ |
| Sex | $F_{(1, 27)} = 18.07; p < 0.0001$ | $F_{(1, 27)} = 16.87; p < 0.0001$ | NS |
| Genotype x Sex Interaction | NS | NS | $F_{(1, 27)} = 5.474; p = 0.0269$ |

Abbreviations: NS, Not significant.

*Post hoc* tests revealed that *App*^NL-G-F^ males have fewer Wake, NREM and REM sleep bouts than WT males across the 24-h period (Fig. 13A**-A****’’**), during the light phase (Fig. 13B**-B****’’**), and for Wake (Fig. 13C) and NREM (Fig. 13**C****’**) during the dark phase. Female *App*^NL-G-F^ mice have fewer REM periods than WT littermate females across all 3 time periods (Fig. 13A**”, B” and C”)** and fewer NREM bouts during the dark phase (Fig. 13**C****’).**

**Figure 13.**
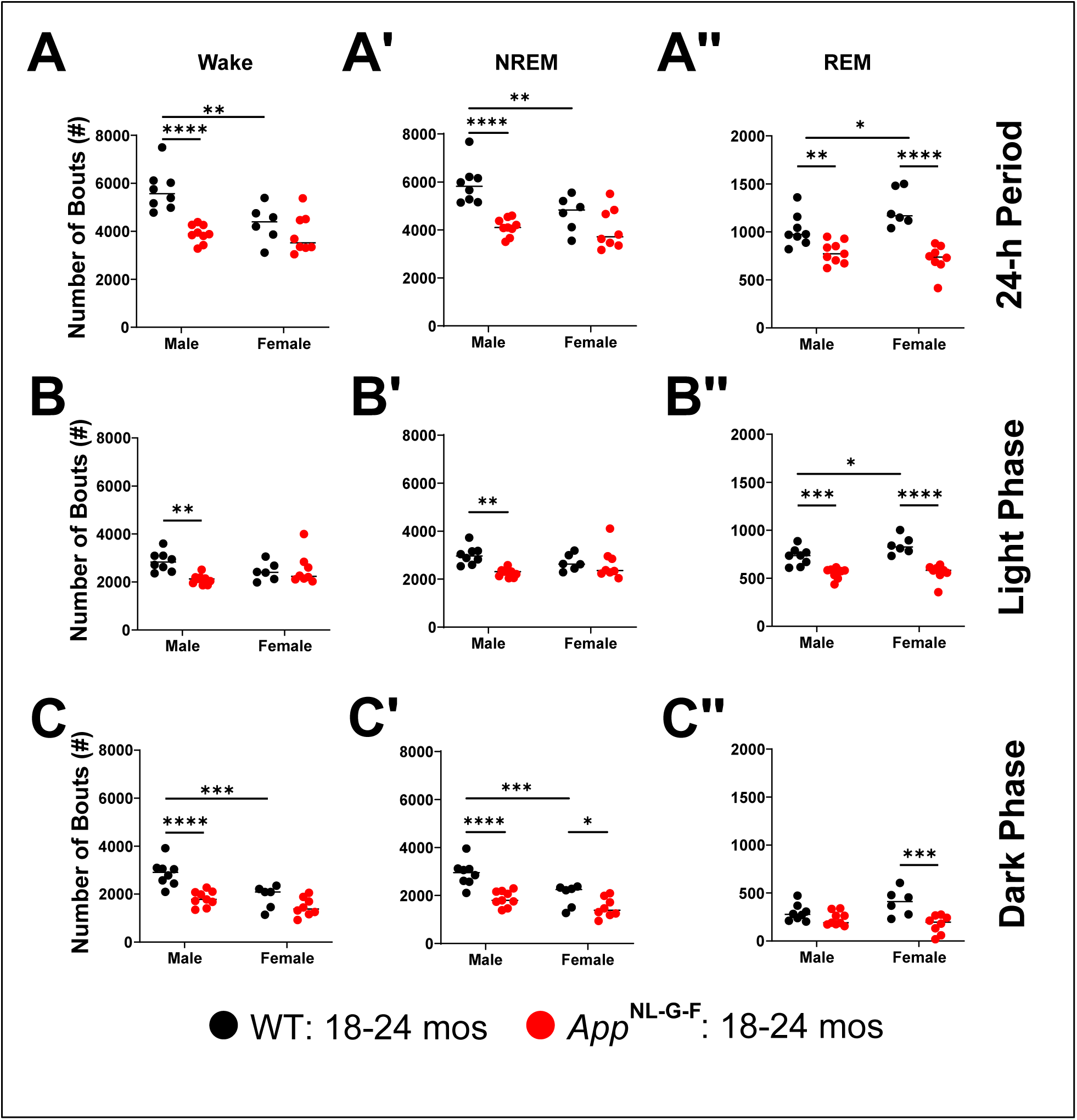
The number of sleep/wake state bouts for male and female *App*^NL-G-F^ and WT mice at 18.3-24.8 months of age. The number of Wake (A-C), NREM (A’-C’), and REM sleep (A’’-C’’) bouts were summed across the 14 24-h periods (A-A’’) recorded as well as the light (B-B’’) and dark (C-C’’) phases. Male *App*^NL-G-F^ mice exhibit fewer Wake (A), NREM (A’) and REM sleep (A’’) bouts in comparison to their age-matched wildtype (WT) littermate controls. This effect is evident in both the light (B-B’’) and dark (C-C’’) phases as well. Female *App*^NL-G-F^ mice exhibit fewer REM sleep bouts than their age-matched wildtype (WT) littermates in all 3 time comparisons. \**p* < 0.05, \*\**p* < 0.01 and \*\*\**p* < 0.001, \*\*\*\**p* < 0.0001 based on between group *post hoc* comparisons using Fisher’s LSD test.

## Discussion

*App*^NL-G-F^ mice at younger ages (6-12 months) have previously been the subject of sleep/wake studies (Calafate et al., 2023; Maezono et al., 2020; Yao et al., 2023). In the present study, we compared the diurnal rhythms of sleep/wake, activity and T_sc_ of *App*^NL-G-F^ mice to WT littermates at 14-18 and 18-22 months of age and evaluated biological sex as a factor in a larger cohort at 18-24 months. In contrast to a single 24-h recording that is typically used to phenotype sleep/wake in most studies including previous studies of *App*^NL-G-F^ mice at younger ages, the sleep/wake data in the older cohort of *App*^NL-G-F^ and *App*^WT^ mice studied here were derived from a continuous 14-day recording. We find that *App*^NL-G-F^ mice have a long wake/short sleep phenotype characterized by extended Wake bouts and reduced REM sleep amounts and, as in human AD patients, that these characteristics are exacerbated in female *App*^NL-G-F^ mice.

### Sleep/wake and activity phenotypes of 14-18 month vs. 18-22 month old App^NL-G-F^ mice

*App*^NL-G-F^ mice exhibited more wake and less NREM and REM sleep than *App*^WT^ mice at both ages (Figure 4). This long wake/short sleep insomnia-like phenotype is also evident at 12 months of age (Maezono et al., 2020) but not at 6 months (Calafate et al., 2023; Maezono et al., 2020), suggesting a progression in this symptomatology. Our observations are consistent with those of Maezono et al. (Maezono et al., 2020) who found reduced REM sleep in *App*^NL-G-F^ mice at both 6 and 12 months of age and increased Wake and reduced NREM sleep at 12 months of age. A prospective study in humans found that lower REM sleep percentage and longer REM sleep latency were both associated with a higher risk of incident dementia (Pase et al., 2017). More recently, a meta-analysis of 24 EEG studies of AD patients found that the most consistent characteristic observed across studies was increased Wake and reduced NREM and REM sleep relative to controls (Y. Zhang et al., 2022), indicating that the long wake/short sleep phenotype of *App*^NL-G-F^ mice resembles that of AD patients.

Although both *App*^NL-G-F^ and WT mice exhibited more activity at 18-22 months than at 14-18 months, increased activity with age was more evident in *App*^NL-G-F^ mice, particularly during the dark phase (Figure 7C), which undoubtedly contributed to the increased amount of wake and reduced amounts of NREM and REM sleep. These changes in sleep/wake and activity were restricted to the dark phase in 14-18 month old mice but occurred during both the light and dark phases at 18-22 months (Figure 5D**, 5D’**). Indeed, *App*^NL-G-F^ mice of both ages showed longer Wake Bout Durations than WT mice during the dark phase; very long Wake Bouts (>260 sec) were significantly longer in *App*^NL-G-F^ than in WT mice (Figure 7A). Overall activity was greater in the older *App*^NL-G-F^ mice during the dark phase (Figure 7C). Whereas previous studies reported that reduced REM sleep occurred during the light phase in 6-12 month old *App*^NL-G-F^ mice (Calafate et al., 2023; Maezono et al., 2020; Yao et al., 2023), we found reduced REM sleep only during the dark phase in both 14-18 month and 18-22 month old *App*^NL-G-F^ mice as described in 9-month old *App*^NL-G-F^ mice (Yao et al., 2023). Irrespective of this difference regarding the timing of reduced REM sleep levels, the results across studies indicate a progressive disruption of the diurnal distribution of sleep/wake that is likely a consequence of the hyperactivity evident in older *App*^NL-G-F^ mice.

### Impact of biological sex on the phenotype of App^NL-G-F^ mice at 18-24 months of age

Robust differences in sleep/wake patterns were present within, as well as between, both WT and *App*^NL-G-F^ strains at 18-24 months of age. Within the wildtype strain, female WT mice had more REM sleep than WT males (**Figure 11A”**) due to more REM sleep bouts during the light phase (**Figure 13B”).** Within the KI strain, female *App*^NL-G-F^ mice had more Wake than males (**Figure 11A**) due to longer Wake bout durations in the dark phase (**Figure 12C**).

Female *App*^NL-G-F^ mice also had less NREM sleep than males (**Figure 11A’**) due to shorter NREM bouts in the light phase (**Figure 12B’**). Sleep/wake patterns in mice are largely determined by strain genetics (Jan et al., 2020; Tafti & Franken, 2002) with minor effects of the estrus cycle (Koehl et al., 2003; Paul et al., 2006; Swift et al., 2024). Our 14-day recordings conducted in 18-24 month old mice (Figure 10) would minimize any effects of the estrus cycle which is typically 4-5 days in mice (Byers et al., 2012).

The within sex/between strain comparisons of sleep/wake characteristics are likely of greater biological interest, particularly whether both sexes of *App*^NL-G-F^ mice exhibit the long wake/short sleep insomnia-like phenotype described above. **Figure 11A** demonstrates that both male and female *App*^NL-G-F^ mice have significantly more Wake than WT mice; this difference was particularly evident in the dark phase (**Figure 11C)** and was due to longer Wake bouts in both sexes at all times of day (Figure 12A-C). Similarly, both sexes of *App*^NL-G-F^ mice exhibited less NREM sleep (**Figure 11A’)** than WT mice; in males, this was due to fewer bouts of NREM sleep (**Figure 13A’)** as described at younger ages (Maezono et al., 2020). Although older male *App*^NL-G-F^ mice show the same tendency toward fewer (**Figure 13A’’)**, longer (**Figure 12A’’)** REM bouts that females do, these differences are less extreme than in females and, consequently, male *App*^NL-G-F^ mice do not exhibit the significant deficit in REM sleep that females do (**Figure 11A’’**).

Until recently (Jin et al., 2025; C. E. Johnson et al., 2024; Y. Zhang et al., 2022), there have been relatively few studies of sleep in AD patients and, despite the fact that AD is more common in women, there is limited information about sex differences in sleep of AD patients. A recent meta-analysis simply stated that “being female and of advanced age were associated with increased SL [sleep latency] in AD patients compared with controls” (Y. Zhang et al., 2022). Women experience sleep disorders at a higher frequency than men including higher rates of insomnia (Zeng et al., 2020; B. Zhang & Wing, 2006) and restless legs syndrome (Berger et al., 2004; Hogl et al., 2005). Sleep apnea rates rise dramatically in women after menopause and both sleep apnea and menopause increase AD risk (Hall et al., 2015; Jehan et al., 2015). CSF biomarkers and PET imaging studies suggest that women with mild cognitive impairment (MCI) and AD exhibit faster brain atrophy rates in both disorders (Barnes et al., 2005; Hua et al., 2010; Jack et al., 2017; Jansen et al., 2015; K. A. Johnson et al., 2016; Mattsson et al., 2017; Shinohara et al., 2016; Skup et al., 2011). Furthermore, women with MCI and elevated amyloid plaque or NFT burden or CSF biomarkers for AD were more likely to develop dementia or neurodegeneration than males, suggesting an elevated susceptibility to AD-related neuropathology in women (Barnes et al., 2005; Koran et al., 2017). Thus, the more severe long wake/short sleep insomnia-like phenotype of female *App*^NL-G-F^ mice resembles some of the symptomatology observed in women with AD.

### Impacts of pathology on sleep/wake and activity

Insomnia in humans is typically characterized by subjective reports of reduced sleep quality and can be accompanied by objective measures of sleep fragmentation and reduced sleep amounts. Both sexes of *App*^NL-G-F^ mice exhibited an insomnia-like phenotype with reduced amounts of both NREM and REM sleep and increased amounts of wakefulness in comparison to WT mice, However, sleep fragmentation was not observed in male *App*^NL-G-F^ mice; paradoxically, these mice exhibited greater sleep consolidation with fewer, longer bouts of both NREM **(Figure 12A’, 13A’)** and REM **(Figure 12A”, 13A”)** sleep, particularly during the light phase **(Figure 12B’, 12B”,13B’, 13B**”). The reduced sleep amounts in male *App*^NL-G-F^ mice was thus due to fewer sleep bouts across the 24-h period. A hyperactive hypocretin/orexin or monoaminergic arousal system or a dysfunctional GABAergic sleep onset system could underlie the longer bouts of Wake in *App*^NL-G-F^ mice (**Figure 12A**). Indeed, Hcrt neurons in aged mice have been shown to exhibit more frequent neuronal activity driving wake bouts and optogenetic stimulation of Hcrt neurons in aged mice results in prolonged wakefulness (Li et al., 2022).

Thus, the hyperactivity evident in Figures 4D**, 4D’,** and **5D’** could drive the longer wake bouts evident in Figure 7A and result in the longer NREM and REM sleep bouts found in male *App*^NL-G-F^ mice (**Figure 12A’ and 12A”**).

Alternatively, Aβ deposition may impair the mechanism(s) that underlies the transition between states such that individual bouts of <u>any</u> state become prolonged. This concept is the converse of that which has been proposed to account for the unstable states that are characteristic of the sleep disorder narcolepsy in which loss of the hypocretin/orexin neurons results in instability of arousal states (Saper et al., 2001; Saper et al., 2010). Impairment of state transitions could either be manifest as a problem to initiate a new state (sleep or wake onset failure) or to terminate the current state.

Female *App*^NL-G-F^ mice also showed fewer REM bouts across the 24-h period (**Figure 13A”**) and longer Wake **(Figure 12A)** and REM **(Figure 12A”)** bouts but, in contrast to males, NREM bout duration was not significantly different **(Figure 12A’)**. The longer Wake and REM bouts occurred in females during both the light and dark phases as well as across the 24-h period (Figure 12). Thus, female *App*^NL-G-F^ mice may share the same impairment of transitions from Wake and REM sleep that male *App*^NL-G-F^ mice exhibit.

Perhaps the most consistently observed characteristic of *App*^NL-G-F^ mice across studies is the deficit of REM sleep (Calafate et al., 2023; Maezono et al., 2020; Yao et al., 2023). As stated above, REM sleep bouts are longer (**Figure 12A”)** and the deficit in REM time is due to many fewer REM bouts in both sexes (**Figure 13A”)**, particularly during the light phase (**Figure 13B”).** This phenotype could result from an impairment in the REM onset circuitry which involves hypocretin and melanin-concentrating hormone (MCH) neurons in the lateral hypothalamus (LH), GABAergic neurons in the ventrolateral periaqueductal grey (vlPAG), glutamatergic neurons in the sublateral dorsal tegmental (SLD) nucleus of the pons, and glycinergic/GABAergic neurons of the ventromedial medulla (Luppi et al., 2025).

Overexpression of the hypocretin/orexin system is often associated with AD and can result in arousals from sleep and impaired REM sleep (Liguori et al., 2014; Musiek & Ju, 2022). A smaller proportion of MCH neurons have been shown to be active during REM sleep in *App*^NL-G-F^ mice and MCH axons in both *App*^NL-G-F^ mice and AD patients exhibit large “spheroid” swellings (Calafate et al., 2023) that disrupt electrical conduction along axons (Yuan et al., 2022).

Moreover, reduced REM sleep in *App*^NL-G-F^ mice has previously been correlated with increased Aβ in the pontine and medullary areas responsible for REM sleep (Maezono et al., 2020). This presumptive impairment of the REM onset mechanism in *App*^NL-G-F^ mice may correspond to the increased REM latency in AD patients (Pase et al., 2017) that has recently associated with higher Aβ burden, phosphorylated tau-181 (p-tau181), and lower brain-derived neurotrophic factor (BDNF) levels (Jin et al., 2025).

### Sleep homeostasis in App^NL-G-F^ mice

Since *App*^NL-G-F^ mice exhibited an insomnia-like phenotype relative to WT mice, we conducted a murine MSLT at 18-22 months of age to determine whether the level of sleepiness differed between the two strains. As indicated in **Figure S1**, there were no significant differences between the two strains in the accumulation of NREM or REM sleep during the five 20 min nap opportunities. We also probed the sleep homeostatic system in these two strains by conducting a 6-h SD beginning at light onset. Both strains increased a conventional measure of sleep homeostasis, EEG Slow Wave Activity, in a comparable manner during the subsequent recovery period (Figure 8E **vs. 8F**). These results differ from those reported in a previous study in which the ratio of EEG delta power recorded during a 4-h rebound after a 4-h SD period relative to the delta power recorded during baseline was reduced in 6 month old *App*^NL-G-F^ compared to WT mice (Calafate et al., 2023). In our study, *App*^NL-G-F^ mice increased the amount of NREM sleep more robustly during the first 2-h of the RS period (Figure 8B) than *App*^WT^ mice (Figure 8A), suggestive of a stronger homeostatic response. When compared across the entire 5-h RS period during the light phase, however, the amount of NREM sleep did not differ between the strains (**Figure S2B).** Together, these results suggest that the amyloid pathology evident in the brains of *App*^NL-G-F^ mice (Figure 2) has minimal impact on the sleep homeostatic system.

### Spectral analysis of the EEG in App^NL-G-F^ mice

During wakefulness, EEG spectral power density declined with age in *App*^NL-G-F^ mice in 4 of the 5 bandwidths and for WT mice in 3 of the 5 bandwidths (Figures 9D-G), indicating an age-related reduction in EEG spectral power in both strains. On the other hand, EEG spectral density in the delta range increased with age for both genotypes, reflecting an overall slowing of the EEG with age (Figure 9C). In contrast to Wake, there were no age-related differences in EEG spectral density during NREM sleep in either strain (**Figure 9C’-H’**). These results differ from the profound reductions in EEG Slow Wave Activity found in NREM of patients with advanced AD (Prinz et al., 1982) and thus suggest that *App*^NL-G-F^ mice may not be a good model of advanced AD. However, EEG spectral power did differ between female *App*^NL-G-F^ and WT mice in the 4.88 −10.50 Hz range during NREM sleep at 18-24 months of age (**Figure 14A**).

During REM sleep, spectral power density in the delta range was greater in *App*^NL-G-F^ compared to WT mice at 18-22 months (**Figure 9C’’**), further indicating a slowing of the EEG with age in *App*^NL-G-F^ mice. Spectral power density was also reduced in the theta band for *App*^NL-G-F^ mice at both ages (**Figure 9D’’**) as occurs in younger *App*^NL-G-F^ mice (Calafate et al., 2023; Maezono et al., 2020). Consistent with previous observations (Maezono et al., 2020), the theta/delta ratio declined in older *App*^NL-G-F^ mice (Figure 9I**”).** Sex-specific analyses revealed reduced EEG spectral power between 6.35 – 11.72 Hz during REM sleep in female *App*^NL-G-F^ mice compared to female WT mice and between 7.08-11.48 Hz for male *App*^NL-G-F^ mice compared to male WT mice (**Figure 14B**). These results are consistent with studies in patients with MCI or early stage AD that report reduced fast oscillatory activity and increased slow oscillatory activity during REM sleep (Brayet et al., 2016; Prinz et al., 1992).

### Consistency of phenotype

In the 18-24 month old cohort, EEG and EMG recordings collected across a 14-day recording period were classified into sleep/wake states using an automated scoring system. This methodology contrasts with the manner in which most sleep/wake studies are conducted, including previous studies of *App*^NL-G-F^ mice (Calafate et al., 2023; Maezono et al., 2020; Yao et al., 2023) as well as our study of Cohort 1. Those studies utilized the traditional sleep phenotyping approach in which EEG/EMG recordings are collected over a single 24-hour period and subsequently analyzed manually. We were interested to know how consistent sleep/wake patterns were over a longer period. While we observed some day-to-day variability in the amounts of wake, NREM, and REM sleep across the 14 days (Figure 10), the overall phenotypes of the 4 sex x genotype groups studied were remarkably consistent and clearly demonstrate that female *App*^NL-G-F^ mice exhibited the most wakefulness and least amount of sleep each day. This approach may allow more robust results to be obtained from smaller cohorts and should be considered as a means to reduce animal use in sleep research.

#### Limitations

*App*^NL-G-F^ mice express humanized *APP,* exhibit a progressive increase in the accumulation of Aβ, a higher ratio of Aβ42 to Aβ40, amyloidosis, and neuroinflammation in several brain areas (Saito et al., 2014) and thus provide numerous advantages over other murine models that overexpress *App*. However, this model only recapitulates some of the AD pathology: tauopathy does not occur and the cognitive phenotype and severe memory deficit associated with clinical onset of AD are not observed in *App*^NL-G-F^ mice (Saito et al., 2014; Sakakibara et al., 2018). Moreover, because studies of this strain used a mutant protein on a powerful exogenous promoter, the effects on sleep described by us and previous investigators may be skewed by overexpression in specific brain areas.

As stated above, *App*^NL-G-F^ mice do not exhibit the profound reductions in EEG Slow Wave Activity found in advanced AD patients (Prinz et al., 1982). As such, *App*^NL-G-F^ mice should be viewed not as a model of advanced AD, but rather as a model of early AD that recapitulates limited pathological components of the disorder. *App*^NL-G-F^ mice may thus provide a translation tool providing insights into the phenotypic changes induced by amyloid pathology, which occurs primarily during the prodromal phase of AD. Neither the present nor previous studies have assessed the effects of age-related changes in brain temperature on sleep/wake, sleep homeostasis or activity.

Analysis of the 14-day EEG recording and classification into sleep stages required the use of automated scoring methods. Although the supervised machine learning algorithms used to segment the data analyzed in the present study are highly accurate, as in other automated sleep/wake methods, the accuracy for identification of REM sleep is lower than for other states. In our previous publications, we have mitigated this limitation by extracting the REM episodes that were most likely to have been misidentified for manual review (Sun et al., 2022; Tisdale et al., 2024). Given the duration of the recordings in the present study, such a manual review was impractical. Our approach instead was to apply rules that we used in our previous studies to identify epochs of REM sleep that were likely to be misidentified arousals or episodes of quiet wakefulness. As shown in **Figure 14B**, spectral analyses of the epochs classified as REM sleep revealed the typical spectral profile of REM sleep with a large peak in the theta frequency range (4-9 Hz), suggesting that our results using Somnivore are reasonable.

#### Conclusions

*App*^NL-G-F^ mice exhibit several characteristics that resemble the sleep phenotype of human AD patients: more wakefulness, less NREM and REM sleep and a slowing of the EEG. However, these mice do not exhibit the reductions in EEG Slow Wave Activity nor the tauopathy and cognitive deficits found in patients with advanced AD. As such, *App*^NL-G-F^ mice may be a better model of the prodromal phase of AD during which sleep interventions may be designed to slow disease progression. The longer Wake bout durations and reduced REM sleep characteristic of App*^NL-G-F^* mice suggest a deficit in the mechanism underlying the transition between states and may be related to the recently-reported delayed REM onset in AD patients. In a fear conditioning paradigm, impaired learning ability has been correlated with REM sleep duration in 13 month old but not 7 month old *App*^NL-G-F^ mice (Maezono et al., 2020). Together, these results support the use of App*^NL-G-F^* mice as a model to investigate sleep-related interventions to mitigate AD burden. Furthermore, these results further support the assertion that sex is an important determinant in disease severity in AD. Specifically, our results suggest that sleep measures, state-specific brain rhythms, and activity are more severely impacted by amyloid pathology in the female sex than in males.

## Materials and Methods

### Animals

Male and female *App*^NL-G-F^ mice that expressed humanized Aβ and the familial Alzheimer’s disease Swedish (K670N, M671L), Arctic (E693G), and Iberian (717, I) mutations on the C57BL/6J background (Saito et al., 2014) were obtained from Drs. Takaomi Saido and Takashi Saito (RIKEN Brain Science Institute, Japan). To obtain wildtype (WT) littermate controls, we crossed heterozygous *App*^NL-G-F/+^ mice to produce homozygous *App*^NL-G-F/NL-G-F^ mice and *App*^+/+^ (WT) littermate controls of both sexes. For simplicity, homozygous *App*^NL-G-F/NL-G-F^ mice are referred to here as *App*^NL-G-F^ mice; heterozygous *App*^NL-G-F/+^ mice were not studied. A total of 31 mice were used in this study, including 17 *App*^NL-G-F^ mice (males: *n* = 9; females: *n* = 8) and 14 age-matched WT littermates (males: *n* = 8; females: *n* = 6). Mice were maintained on a LD12:12 light:dark cycle at room temperature (22 ± 2°C; 50 ± 20% relative humidity) and had access to food and water *ad libitum*. Due to hardware limitations, mice were studied in two cohorts of N=16. Each cohort was initially balanced at N=4 with respect to sex and genotype but became imbalanced over time due to mortality and subsequent substitution of replacement mice. The experimental procedures used here were approved by the Institutional Animal Care and Use Committees at SRI International and at the Gladstone Institute.

### Surgical procedures

Mice were anesthetized with isoflurane and sterile telemetry transmitters (HD-X02, Data Sciences International., St Paul, MN) were placed subcutaneously on the left dorsum. At the time of surgery, *App*^NL-G-F^ mice from Cohort 1 ranged in age from 14.0-17.8 mos (16.0 ± 0.5 mos) and WT mice were from 14.3-17.9 mos (15.6±0.6 mos) whereas Cohort 2 *App*^NL-G-F^ mice ranged from 20.1-23.4 mos (21.7±0.4 mos) and WT littermates were 20.2-23.4 mos (22.0±0.4 mos). Biopotential leads were routed subcutaneously to the head and EMG leads were positioned in the right nuchal muscle. Cranial holes were drilled through the skull at −2.0 mm AP from bregma and 2.0 mm ML at −1.0 mm AP and −1.0 mm ML from lambda. The two biopotential leads used as EEG electrodes were inserted into these holes and affixed to the skull using dental acrylic. The placement of HD-X02 telemetry transmitters also allowed measurement of subcutaneous body temperature (T_sc_) as well as locomotor activity. The incision was closed with absorbable suture. Analgesia was managed with ketoprofen (5 mg/kg, s.c.) and buprenorphine (0.05 mg/kg, s.c.) upon emergence from anesthesia and for the first day post-surgery. Meloxicam (5 mg/kg, s.c., q.d.) was continued for 2 d post-surgery. Mice were singly housed post-surgery and for the duration of the experiment.

### Experimental design

Mice were recorded in a general mouse housing room at the Gladstone Institutes. DSI receiver boards were placed within two conventional animal housing racks with 8 receiver boards per rack in which each rack accommodated 16 cages, front and back. The singly-housed mice were thus recorded surrounded by cages in which there were usually multiple mice per cage. Food and water were available *ad libitum* and lighting was maintained on a 12:12 light/dark cycle in a temperature- and humidity-controlled environment.

Experiment 1 – Longitudinal comparison of *App*^NL-G-F^ vs. WT mice at 14-18 and 18-22 <u>months of age</u>. Cohort 1 mice underwent the sequence of procedures illustrated in Figure 1A. Beginning no sooner than 1 week post-surgery when the mice were 14.5-18.2 months of age (*App*^NL-G-F^: 16.3 ± 0.5 mos; WT: 15.9 ± 0.6 mos), EEG and EMG were continuously recorded for 14 days. Mice were again recorded continuously for another 14 days when they ranged in age from 18-22 (*App*^NL-G-F^: 20.4 ± 0.5 mos; WT: 19.8 ± 0.7) months. Due to mortality of one female *App*^NL-G-F^ and one female WT mouse in the intervening 16 weeks between recordings, a male *App*^NL-G-F^ mouse of comparable age was substituted and was recorded only at the older age.

Upon conclusion of the second 14-day recording, Cohort 1 mice underwent a murine Multiple Sleep Latency Test (MSLT) to evaluate their basal level of sleepiness (Veasey et al., 2004). During the following week, Cohort 1 mice underwent a 6-h sleep deprivation (SD) beginning at light onset (ZT0) followed by an 18-h EEG/EMG recording to determine the response to perturbation of sleep homeostasis. Six weeks later, when the remaining 8 mice in Cohort 1 ranged from 20-24 months of age, they were deeply anesthetized, perfused with heparinized saline and 4% paraformaldehyde, post-fixed and frozen for subsequent histology along with 6 age-matched males (4 *App*^NL-G-F^; 2 WT) that had not been implanted.

Experiment 2 – Comparison of *App*^NL-G-F^ vs. WT mice at 21-24 months of age. Cohort 2 mice underwent a 14-day continuous EEG/EMG recording when they ranged in age from 21.0-24.3 mos (*App*^NL-G-F^: 22.3 ± 0.4 mos, N=8; WT: 22.8 ± 0.4 mos, N=8) to complement the data collected for Cohort 1 at 18-22 months (Figure 1B). At the conclusion of the 14-day recording, all 16 mice were perfused and brains removed for subsequent histology. For Experiment 2, the data collected from Cohort 1 at 18-22 months were combined with the data collected from Cohort 2 at 21.0-24.3 months so that analyses were conducted on N=31 mice as described in *Animals* above.

### Classification of arousal states

EEG/EMG recordings were scored in 10-s epochs as either Wake, non-Rapid Eye Movement (NREM) or REM sleep by experienced scorers using Somnivore (Allocca et al., 2019). Wake was determined by mixed frequencies in the EEG and a relatively high muscle tone measured in the EMG. NREM sleep was identified by low frequency, high amplitude activity in the EEG and moderate EMG levels. REM sleep was scored when the EEG exhibited low to moderate amplitude, mixed frequency activity accompanied by low muscle tone in the EMG. During REM sleep, theta activity was often the predominant frequency in the EEG and phasic activity was often observed in the EMG. The 14-day recording periods for each mouse were provided to Somnivore (version 1.1.7.0) in EDF format, from which 100 10-s epochs of each state were selected by an expert human scorer as training data. The selected data were used to train a classifier for each mouse and applied in a supervised machine learning model to automatically score each state for the duration of each recording. Following this initial scoring, the accuracy of the 3-4 autoscored hour-long bins was assessed visually. If the automated scoring was determined by the expert human scorer not to generalize well, additional epochs were provided to the training dataset and automatic scoring of the recording was repeated. The following scoring rules were applied to each recording: (1) single epochs of REM sleep were rescored as the same state as the previous epoch (i.e., a minimum of two consecutive 10-s epochs were required for a bout of REM sleep to be scored); and (2) REM sleep directly following wake was rescored as wake (i.e., sleep onset REM periods were excluded). For the MSLT recordings conducted in Cohort 1 mice, the same classifier used to score the baseline recordings was implemented, and manual review and adjustment of the MSLT period was performed.

### Data Analysis and Statistics

<u>General Procedures</u>. In addition to the time spent in each state, the duration and the number of bouts for each state were determined. A “bout” of a particular state was defined as two or more consecutive epochs of that state and ended with a single epoch of any other state.

The EEG power spectrum (0.5–60 Hz) during NREM and REM sleep was analyzed by fast Fourier transform algorithm on all epochs of each state. For spectral analyses, the first 2 transition epochs of each state were excluded. EEG spectra for each state were analyzed in 0.061 Hz bins and in standard frequency bands rounded to the nearest half-Hz value (delta: 0.5–4 Hz, theta: 6–9 Hz, alpha: 9–12 Hz, beta: 12–30 Hz and low gamma: 30–60 Hz). Sleep/wake state amounts and sleep/wake architecture measures (bout durations, number of bouts) were analyzed by ANOVA as described below. Statistics were calculated using GraphPad Prism (ver. 10.0.2). For the EEG spectra data, statistical comparisons for standard frequency bands were conducted as described for sleep architecture measures. Additionally, unpaired t-tests with corrections for multiple comparisons using the Holm-Šídák method were performed for each 0.061 Hz bin from 0.5-60 Hz between genotypes and within each sex.

<u>Experiment 1 data analysis</u>. As stated above, EEG/EMG recordings were conducted in a general mouse housing room at the Gladstone Institute. To obtain relatively undisturbed representative recordings of baseline sleep/wake in Cohort 1 *App*^NL-G-F^ vs. WT mice, 24-h periods recorded on a Sunday during the 14-day data capture when the mice were 14-18 months of age and 4 months later when Cohort 1 mice were 18-22 months old were selected for analysis and comparison between strains. After the recordings were scored as described above, a mixed-effects model analysis of variance (ANOVA) was conducted with Genotype, Age, and Time of Day as factors. When ANOVA indicated statistical significance, Fisher’s least significant difference (LSD) tests were performed *post hoc* to determine specific differences.

<u>Experiment 2 data analysis</u>. Recordings collected throughout the entire 14-day period when Cohort 2 *App*^NL-G-F^ and WT mice were 21.0-24.3 months of age were scored, combined with the data from the recordings of Cohort 1 mice at 18-22 months of age, and the combined data from the *App*^NL-G-F^ (N=17) and WT (N=14) strains were then compared. Data were analyzed as time spent in each state as well as during the dark and light phases across the recording period. The larger sample size in Experiment 2 (N=31 mice) allowed a mixed-effects model ANOVA to be conducted with Genotype, Sex, and Time of Day as factors followed by between group *post hoc* comparisons using Fisher’s LSD when appropriate.

### Histology and immunohistochemistry

*App*^NL-G-F^ mice were anesthetized and transcardially perfused with 0.1 M phosphate buffer saline (PBS), and brains were extracted and post-fixed in 4% phosphate-buffered paraformaldehyde at 4°C for 48 h. After rinsing with PBS, brains were transferred to 30% sucrose in PBS at 4°C for 24 h and coronally sectioned with a sliding microtome. Ten subseries of floating sections (30 µm) were collected per mouse and kept at 20°C in cryoprotectant medium until use. Each subseries contained sections throughout the rostrocaudal extent of the forebrain. Brain sections were washed 3x for 10 min with PBS to remove cryoprotectant medium and once with 0.5% Tween-20 (PBS-T) to permeabilize the tissue. Endogenous peroxidases were blocked with a 15-min incubation of 3% hydrogen peroxide and 10% methanol in PBS. Sections were subsequently washed 3x for 10 min each in PBS-T. Nonspecific binding was blocked with a blocking solution containing 10% normal donkey serum (Jackson ImmunoResearch, 017-000-121) and 0.2% gelatin (Sigma-Aldrich, G2500) in PBS-T for 1 h. Brain sections were incubated overnight with biotinylated 82E1 mouse anti-Aβ antibody (IBL, 10326) at 1:500 and 1:1000 dilution for rabbit anti-Iba1 antibody (Wako, 019–19741) in 3% normal donkey serum 0.2% gelatin and PBS-T at 4°C. For anti-Iba1 staining, after washes, brain sections were incubated with biotinylated donkey anti-mouse antibody (Jackson ImmunoResearch) at 1:500 dilution for 2.5 h at room temperature (RT). Unbound antibodies were removed with three washes of 10 min with 0.5% PBS-T and one wash with PBS for 10 min. Iba1 and 82E1 stainings were developed with an avidin-biotin complex (ABC) kit (Vector Laboratories, PK-6100) according to the manufacturer’s protocol and sections were incubated for 1 h at RT. Brain sections were subsequently washed three times for 10 min each with PBS at RT and incubated with diaminobenzidine (Vector Labs) for colorimetric development.

Sections were then washed 3x for 10 min each with PBS and mounted in 1.2% gelatin in H_2_O and allowed to dry. After two 5-min washes in xylene, brain sections were permanently mounted on slides and coverslipped for analysis.

### Quantification of immunostaining

DAB-stained sections (82E1 and Iba1) were imaged using a Leica Versa 200 Slide Scanner under identical exposure settings across samples. Area coverage (%) for plaque burden (82E1) and microglial coverage (Iba1) was quantified using ImageJ. Regions of interest included the entire cerebral cortex (CTX), retrosplenial cortex (RSC), posterior parietal cortex (PPC), somatosensory cortex (SSC), and hippocampus (HPC), which were manually outlined based on anatomical boundaries defined by the Allen Mouse Brain Atlas. For each region, stained signal was auto-thresholded, binarized, and measured, and values are reported as percent area coverage relative to the total region area. Due to variation in Iba1 immunostaining across WT mice, Iba1 measurements were normalized to WT such that the mean Iba1 area coverage for WT mice within each region of interest was set to 1. The negligible 82E1 signal in WT mice obviated the need for normalization.

## Acknowledgements

We thank Takashi Saito of the Nagoya University Graduate School of Medical Science and Takaomi Saido of the RIKEN Center for Brain Science for providing the mice and Jasmine Heu and Charlene Sykes of SRI International for providing technical support.

## Author Contributions

RKT, SRM, JJP and TSK designed the study. RKT, SRM, SML and JS executed the study.

Data collection occurred in laboratory space assigned to JJP at the Gladstone Institute. YS, SP, JS and TSK analyzed the results. GA provided analytical software. RKT, YS, JS and TSK wrote the manuscript.

## Data Availability

Upon acceptance of this manuscript for publication, all data will be deposited at the National Sleep Research Resource (https://sleepdata.org) as we have done previously.

## SUPPLEMENTARY INFORMATION

**Figure S1.**
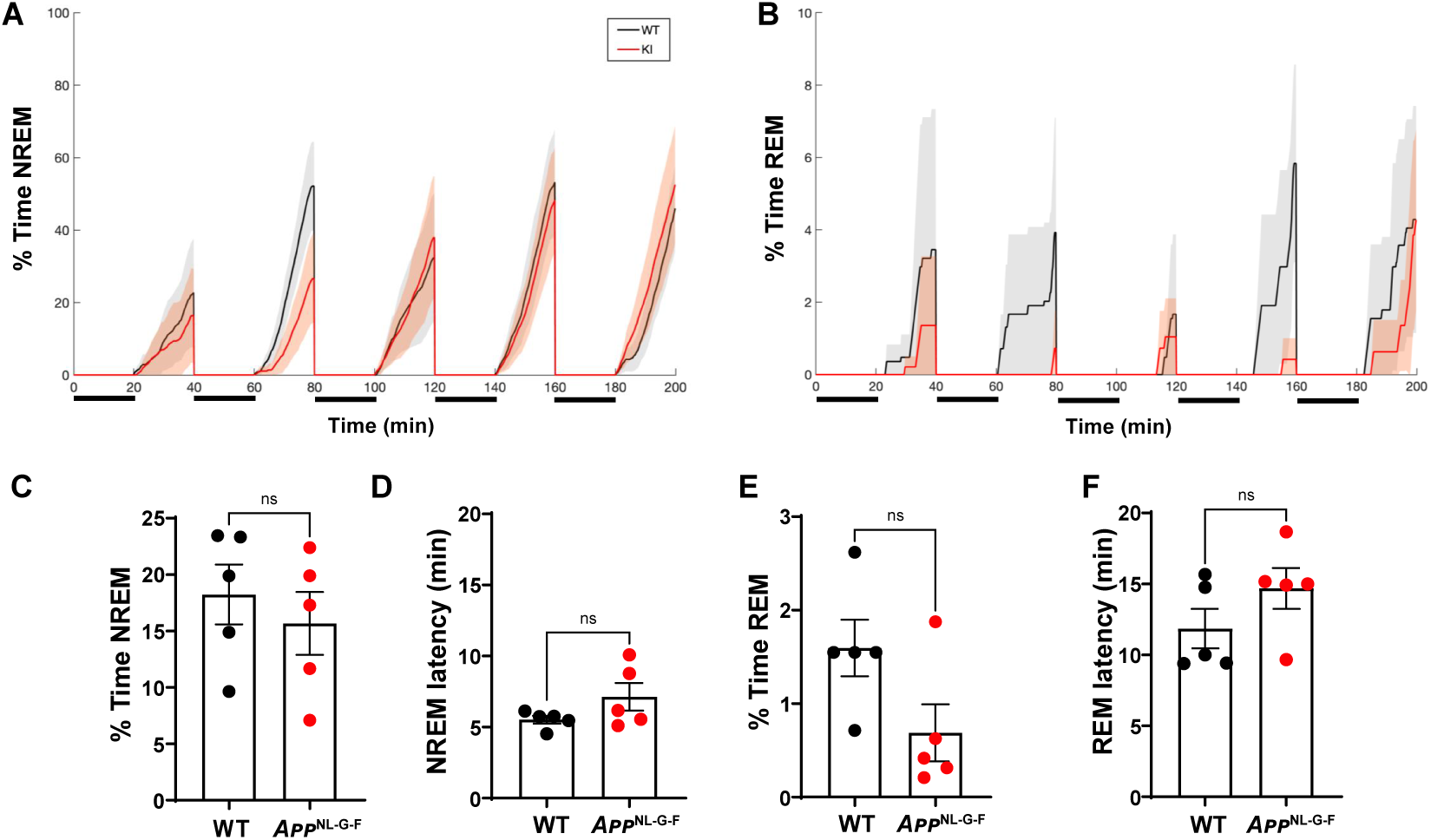
Multiple Sleep Latency Test (MSLT) results from 18-24 month old *App*^WT/WT^ (WT) and *App*^NL-G-F^ mice. **A**. Percent time in NREM sleep during each 20 min nap opportunity in WT (black) and *App*^NL-G-F^ (red) mice. The five 20 min SD periods are indicated by horizontal bars below the abscissa. **B.** Percent time in REM sleep during each 20 min nap opportunity. **C.** Mean percent NREM time during the 5 nap opportunities for WT and *App*^NL-G-F^ mice. **D.** Mean NREM sleep latency in the WT and *App*^NL-G-F^ mice. **E**. Mean percent REM time during the 5 nap opportunities in WT and *App*^NL-G-F^ mice. **F**. Mean REM sleep latency between WT and *App*^NL-G-F^ mice. Values are mean ± SEM. **, *p* < 0.01.

**Figure S2.**
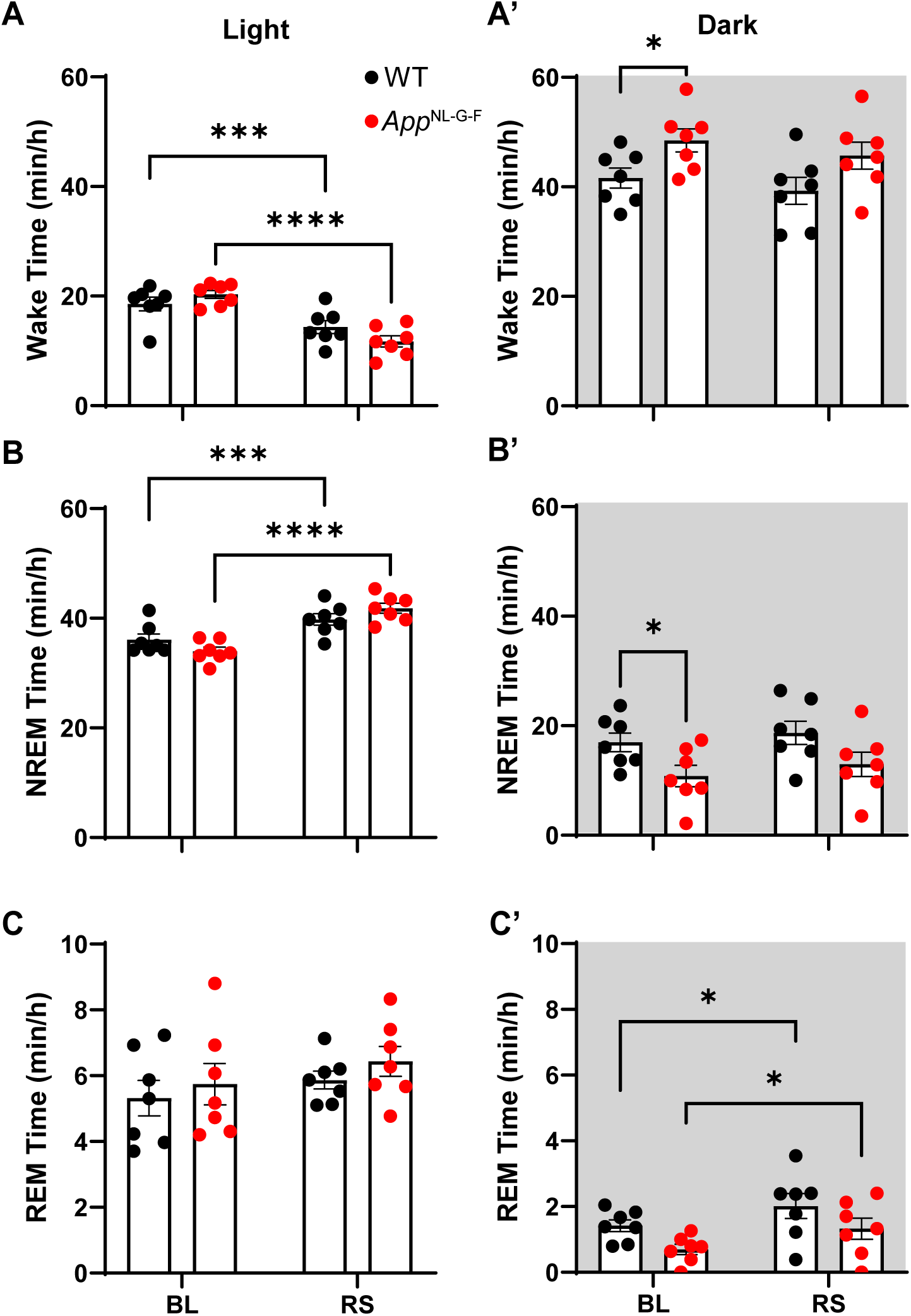
Arousal state amounts during baseline (BL) and during recovery sleep (RS) during the 6-h of the light phase after cessation of 6-h sleep deprivation (left) and the subsequent 12-h dark phase (right) in *App*^WT/WT^ (WT) and *App*^NL-G-F^ mice. **A** and **A’**. Mean hourly amounts of Wakefulness. **B** and **B’**. Mean hourly amounts of NREM sleep. **C** and **C’.** Mean hourly amounts of REM sleep. Values are mean ± SEM. *, *p* < 0.05; **, *p* < 0.01; ***, *p* < 0.001; ****, *p* < 0.0001.

